# Categorical representations of decision-variables in orbitofrontal cortex

**DOI:** 10.1101/135707

**Authors:** Junya Hirokawa, Alex Vaughan, Adam Kepecs

**Affiliations:** Cold Spring Harbor Laboratory, 1 Bungtown Road, Cold Spring Harbor, NY, 11724; Current address: Doshisha University, Kyotanabe, Kyoto 610-0394, Japan

## Abstract

The brain creates internal representations of the external world in the form of neural activity, which is structured to support adaptive behavior. In many cortical regions, individual neurons respond to specific features that are matched to the function of each region and statistics of the world. In frontal cortex, however, neurons display baffling complexity, responding to a mixture of sensory, motor and other variables. Here we use an integrated new approach to understanding the architecture of higher-order cortical representations, and use this approach to show that discrete groups of orbitofrontal cortex (OFC) neurons encode distinct decision variables. Using rats engaged in a complex task combining perceptual and value guided decisions, we found that OFC neurons can be grouped into distinct, categorical response types. These categorical representations map directly onto decision-variables of a choice model explaining our behavioral data, such as reward size, decision confidence and integrated value. We propose that, like sensory neurons, frontal neurons form a sparse and over complete population representation aligned to the natural statistics of the world – in this case spanning the space of decision-variables required for optimal behavior.

## Introduction

Principles of neural representation have been identified in many brain areas by analyzing which features of the external environment activate individual neurons (DiCarlo et al., 2012; Frank et al., 2000; Vinje and Gallant, 2000). For example: in visual cortex, neurons respond to edges; in hippocampus, an area involved in episodic memories, neurons respond in specific spatial and temporal context. When examining frontal areas engaged in decision-making, however, one is struck most of all by the complexity and diversity of their neuronal responses, and similar general principles have been elusive.

Neurons in frontal cortex typically respond to a complex set of variables, and encode a mixture of stimulus features, task parameters, and internal variables (Brody, 2003; Machens et al., 2010; Mante et al., 2013; Rigotti et al., 2013; Wallis and Miller, 2003) This property is known as *mixed selectivity* (Machens et al., 2010; Mante et al., 2013; Rigotti et al., 2013). Within OFC, individual neurons appear to encode key variables such as *outcome expectancy* (expectation of a reward), *reward risk* (outcome variance), *decision confidence* (estimated probability that a decision is correct), or *chosen value* (the overall utility of a choice) (Abe and Lee, 2011; Farovik et al., 2015; Feierstein et al., 2006; Kennerley et al., 2011, 2009; Kepecs et al., 2008; McGinty et al., 2016; Morrison and Salzman, 2009; O’Neill and Schultz, 2010; Ogawa et al., 2013; Padoa-Schioppa and Assad, 2006; Padoa-Schioppa, 2013; Roitman and Roitman, 2010; Schoenbaum et al., 1998; Steiner and Redish, 2014; Sul et al., 2010; Thorpe et al., 1983; Tremblay and Schultz, 1999; Wallis and Miller, 2003). These single-neuron results can be difficult to generalize, because of the apparent diversity in neural response types. One approach to understanding responses across the full neural population, which we refer to as the “model-based” framework, seeks to decipher cortical populations through computational models of behavior. After formalizing cognitive concepts such as “reward value” or “weight of evidence” as quantitative decision-variables that explain animal behavior, these decision-variables can be “decoded” from the population neuronal activity using regression models (Gold and Shadlen, 2000; Lee, D., Seo H. et al., 2012; Platt and Glimcher, 1999). This approach has shown that a variety of decision-variables are well-represented in frontal populations (Gold and Shadlen, 2007; Gutierrez et al., 2006; Kiani et al., 2014; Rich and Wallis, 2016; Rushworth et al., 2011; Schoenbaum and Eichenbaum, 1995; van Duuren et al., 2008).

An alternate, “model-free” framework, embraces the diverse selectivity of cortical responses, since it can increase computational capacity (Sussillo and Abbott, 2009), provide for robust attractor dynamics (Mante et al., 2013; Rigotti et al., 2010) and support learning of complex nonlinear associations (Rigotti et al., 2013). This framework is supported by the observation of *random mixed selectivity* in some populations, in which the precise mixture of features encoded by each neuron are chosen independently (Pagan and Rust, 2014; Raposo et al., 2014).

However, neither framework explains how neuronal populations are structured to support behavior. While model-based decoding is powerful, a range of related decision-variables can often be decoded with equal plausibility – especially if *random mixed selectivity* holds. In contrast, the model-free framework does not provide insight into the single neuron representations of decision variables such as reward, confidence, or value.

Here, we present an integrative approach to understand neural representations in higher-order areas that bridges these frameworks. Using a *model-free* analysis of OFC neurons we show that, despite overwhelming diversity, the population as a whole can be classified into a small set of categorical representations. Using a *model-based* approach, we then map these categorical representations directly onto quantitative decision variables that appear to drive behavior in our task, such as expected reward, decision confidence, and choice value. This approach synthesizes our understanding of single neuron and population representations, and provides a general model of cortical representation as an over-complete dictionary that supports optimal behavior.

## RESULTS

To engage orbitofrontal cortex, we designed a task that would allow a characterization of neural response properties across a large behavioral space. We trained rats to perform a reward-biased, two-choice olfactory discrimination task, in which animals must determine which of two odors dominates in an olfactory mixture (Fig. 1a). Choices to the response port associated with the dominant odor are rewarded with water after a 1s delay, while incorrect choices elicit white noise after 1s without reward. In this task, perceptual uncertainty is manipulated by varying the ratio of odors in the mixture between trials, while reward expectations are manipulated by varying reward size for each port across blocks of trials (Fig. 1b). For each rat, choice accuracy varied systematically with stimulus difficulty, ranging from nearly perfect (pure odors) to near chance (difficult odor mixtures) (Fig. 1c, left). Block-wise changes in reward size induced rapid and sustained choice biases (5.45 ± 0.48 trials to zero-bias after block transitions, mean ± SEM) (Fig. 1d) and motivational shifts that were reflected in reaction times (Fig. 1c, right)(Zariwala et al., 2013). Indeed, the animals showed choice bias that was proportional to stimulus difficulty (Fig. 1c, center) and acquired greater total reward in each session than expected from naive strategies based on odor stimuli or reward size alone (Extended Data Fig. 1), suggesting optimal integration of perceptual uncertainty and reward history.

**Figure 1:**
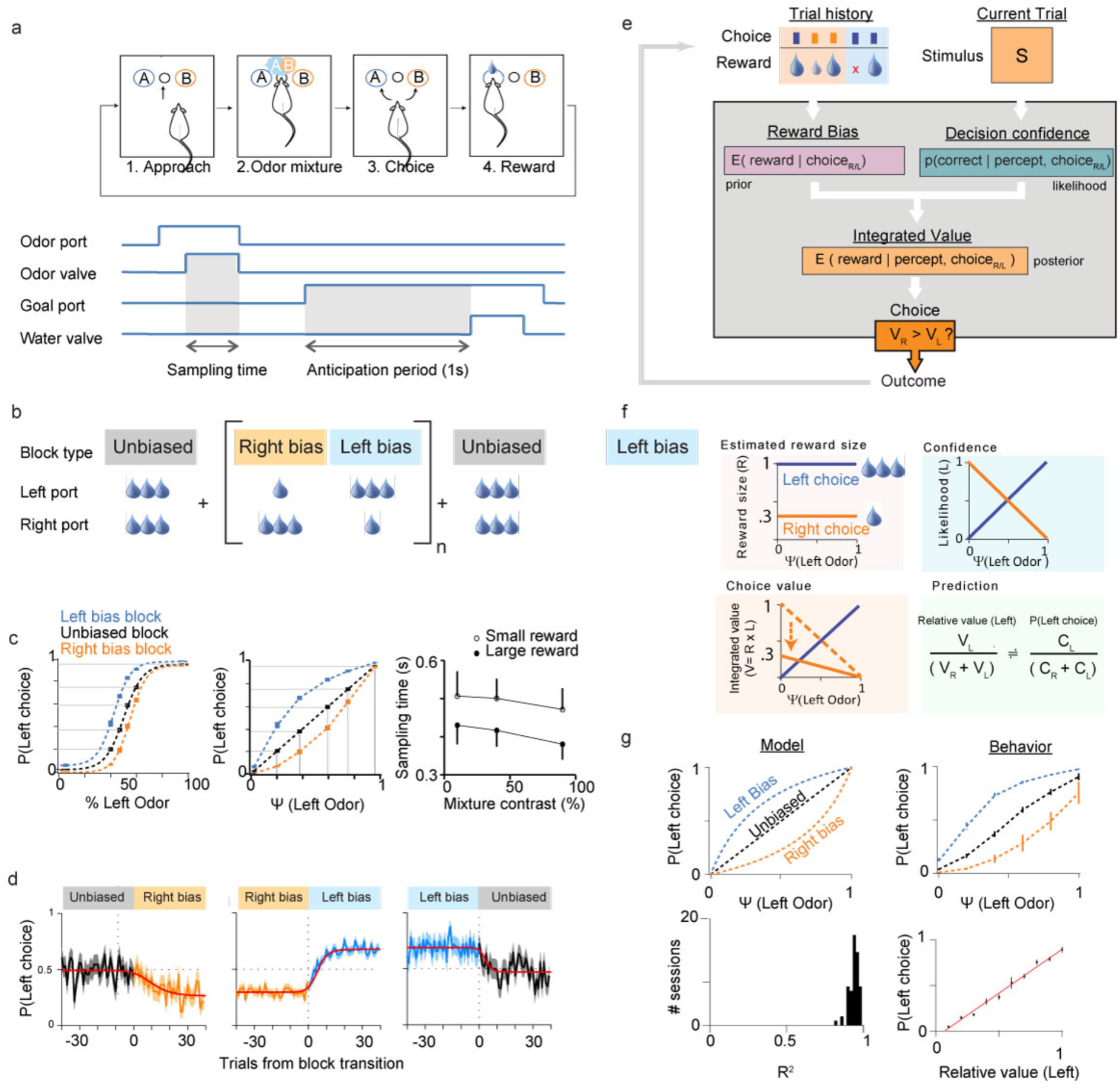
Reward-biased psychometric odor discrimination task. a. Task design for a single trial. **b** Design of block-wise reward manipulation across a behavioral session. A session begins with a control block (unbiased reward) followed by 3 sets of alternating left/right biased blocks, and ends with unbiased block. **c** Psychometric response (*left*) Averaged psychometric functions in control block, left bias block, and right bias block from three rats. (*center*) Linearization of stimuli. For each session, odor stimuli were varied to elicit a consistent range of choice probabilities for the unbiased blocks; these choice probabilities are used to generate a linearized response function Ψ(Left Odor) that is utilized as an internal variable to explain choice probability in odor-based decisions. *(right)* For all animals, odor sampling time was larger for small rewards than large rewards. ***d*** Choice biases were acquired rapidly across block changes (mean +/- SEM) (*right*). Average transition curve (red) was fitted by logistic nonlinear regression model (see Methods). ***e*** Decision-variable model of animal behavior. This model integrates variables from trial history (choice, size and presence of reward) and the current stimulus to generate internal variables representing *reward bias* and *decision-confidence*. These are integrated to generate an estimate of *choice value*. ***f*** Variables arising from the decision-variable model include reward size, confidence, and choice value. The model approximates animals’ choice probability with relative values. ***g*** The decision-variable model accurately predicts (*top left*) changes in choice probability arising due to reward biases (*top right*). The model provides an accurate fit across sessions (*bottom left)* driven by the relative value of each choice (*bottom right*).

To understand the behavioral strategies employed, we first used a model-based model to examine whether animals could optimally integrate decision-variables derived from reward history and sensory stimuli (Fig 1e-f). We assumed that animals combine three distinct decision-variables to evaluate each choice: (i) *decision confidence,* the trial-by-trial estimated probability that each choice is correct based on the evidence, (ii) *reward size,* a block-wise estimate of the relative reward size associated with each choice, and (iii) *previous outcome*, the presence or absence of reward in the preceding trial. Integrating these variables in a multiplicative way, our model provides a normative prediction for the overall value of a choice, termed *choice value.* As evidence that animals use this model, we note that animals outperform the simpler model lacking integration of evidence or reward size (Extended Data Fig. 1). Indeed, this estimate of *choice value* explains two key features of animal behavior: biases arising from block-wise changes in reward size (Fig 1g) and trial-by-trial biases arising from the outcome of previous trials (Extended Data Fig. 2 a-d). These results suggest that animals optimize their choice strategy to maximize total reward by combining decision confidence and reward values, and that a model encoding these decision-variables supports the major features of animal behavior in this complex task.

**Table 1.**

|  | <u>Pooled</u> |  | <u>Rat C068</u> |  | <u>Rat C051</u> |  | <u>Rat C052</u> |  | Chi-square test w/<br>Bonferroni correction |
| --- | --- | --- | --- | --- | --- | --- | --- | --- | --- |
|  | Neurons | % | Neurons | % | Neurons | % | Neurons | % |  |
| Total | 485 |  | 268 |  | 152 |  | 65 |  |  |
| Cluster 1 | 100 | 21% | 64 | 24% | 23 | 15% | 13 | 20% | p = 0.74 |
| Cluster 2 | 97 | 20% | 57 | 21% | 31 | 20% | 9 | 14% | 1.00 |
| Cluster 3 | 61 | 13% | 19 | 7% | 23 | 15% | 19 | 29% | 0.00 * |
| Cluster 4 | 57 | 12% | 28 | 10% | 15 | 10% | 14 | 22% | 0.21 |
| Cluster 5 | 49 | 10% | 22 | 8% | 22 | 14% | 5 | 8% | 0.55 |
| Cluster 6 | 42 | 9% | 24 | 9% | 16 | 11% | 2 | 3% | 1.00 |
| Cluster 7 | 35 | 7% | 24 | 9% | 11 | 7% | 0 | 0% | 0.25 |
| Cluster 8 | 23 | 5% | 10 | 4% | 10 | 7% | 3 | 5% | 1.00 |
| Cluster 9 | 21 | 4% | 20 | 7% | 1 | 1% | 0 | 0% | 0.01 * |

To explore the information present in orbitofrontal cortex, we recorded 485 well isolated neurons from the lateral OFC in 3 rats performing this task (Extended Data Fig. 3). We focused our analysis on the anticipation period, when animals fixate their nose in a choice port awaiting an uncertain reward (Fig. 1a). In this epoch reward prediction may be examined without potential confounds from the sensory stimulus, choice, motor activity, or knowledge about the upcoming reward. First, we sought to identify individual OFC neurons that encode the specific variables present in our model, i.e., decision confidence, reward size, and chosen value (Extended Data Fig. 4). As expected, we observed neurons with response profiles consistent with a representation of decision confidence (Fig. 2a-d) (Hangya et al., 2016; Kepecs et al., 2008). In the anticipation period, this neuron shows ramping activity that is proportional to the evidence supporting each choice (Fig. 2a). Importantly, this neuron was not influenced by block-wise variations in expected reward size – despite the strong influence of reward bias on choice behavior (Fig. 2b-d). We also observed neurons encoding reward size, i.e., anticipated reward assuming that a given choice is correct (Fig. 2e-f). During bias blocks, these neurons showed maximal activity when the animal has chosen the port associated with a small reward. Lastly, we observed neurons representing the overall value of the chosen option, which is proportional to both the probability of reward (i.e. decision confidence in this task) and the expected reward size (Fig. 2g-h).

**Figure 2.**
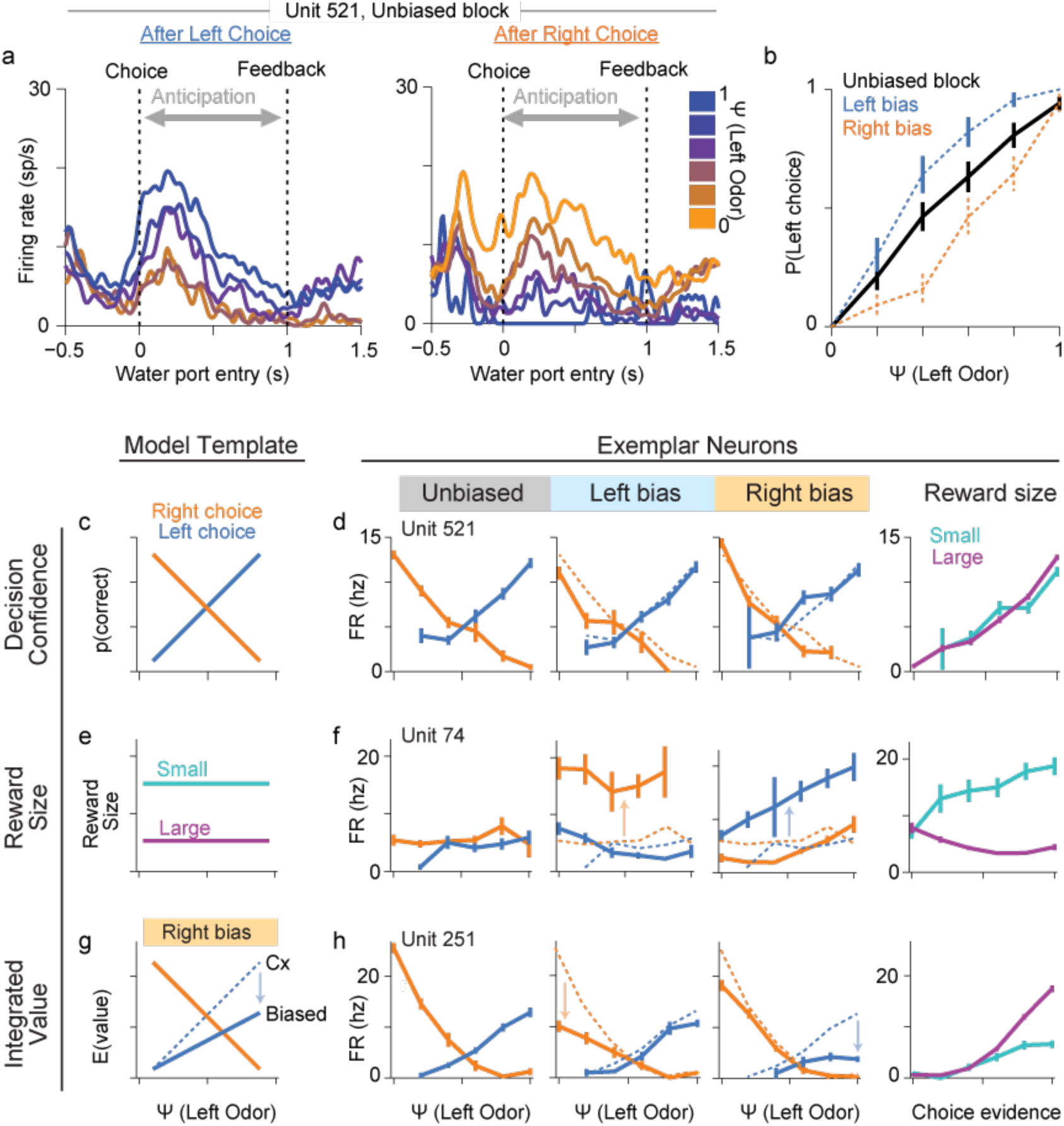
Example neurons from OFC. *a*. Peri-event response histogram for Unit 521 recorded during a control block. Firing rate during the anticipation window was highest when the stimulus supported the choice that was made. Neural activity is aligned to the timing of entry into the choice port. Water reward or an error tone was delivered 1s after entry into the choice port. ***b*** During the recording session for Unit 521, the animal showed significant behavioral bias towards larger rewards. ***c-d*** The template for a representation of decision confidence (**c**) is well matched by the response profile of Unit 521 (**d**). The response profile is generated from averaged activity across the 1s anticipation window for each trial condition – after a choice has been made, but before reward is delivered. This neuron shows a strong response after a choice supported by strong stimulus evidence, but no influence of expected reward size. ***e-f*** The template for a representation of expected reward size (**e**) is well matched by Unit 74. The response of this neuron is not influenced by the odor stimulus, but shows a strong expectancy signal for the upcoming reward. ***g-h*** The template for a representation of integrated value (**g**) is well matched by Unit 251. This neuron is driven by a combination of odor stimulus, choice, and expected reward size that varies across bias blocks type.

For analysis of the entire population, we generated behavioral tuning curves for all 485 OFC neurons by computing their mean activity during the anticipation epoch across all contingencies: stimuli (6 odor mixtures), reward blocks (3 types), choice (left/right), and the outcome of the previous trial (correct/error). After dropping rare conditions, this representation yielded a 42-dimensional behavioral tuning vector for each neuron (Fig. 3a-b), which was de-noised using probabilistic PCA to retain 90% variance (21 dimensions, Extended Data Fig. 5b). Examination of the major eigenvectors reveals that many decision-variable representations are strongly represented and highly variable across the OFC population (Fig. 3b and Extended Data Fig. 5a).

**Figure 3.**
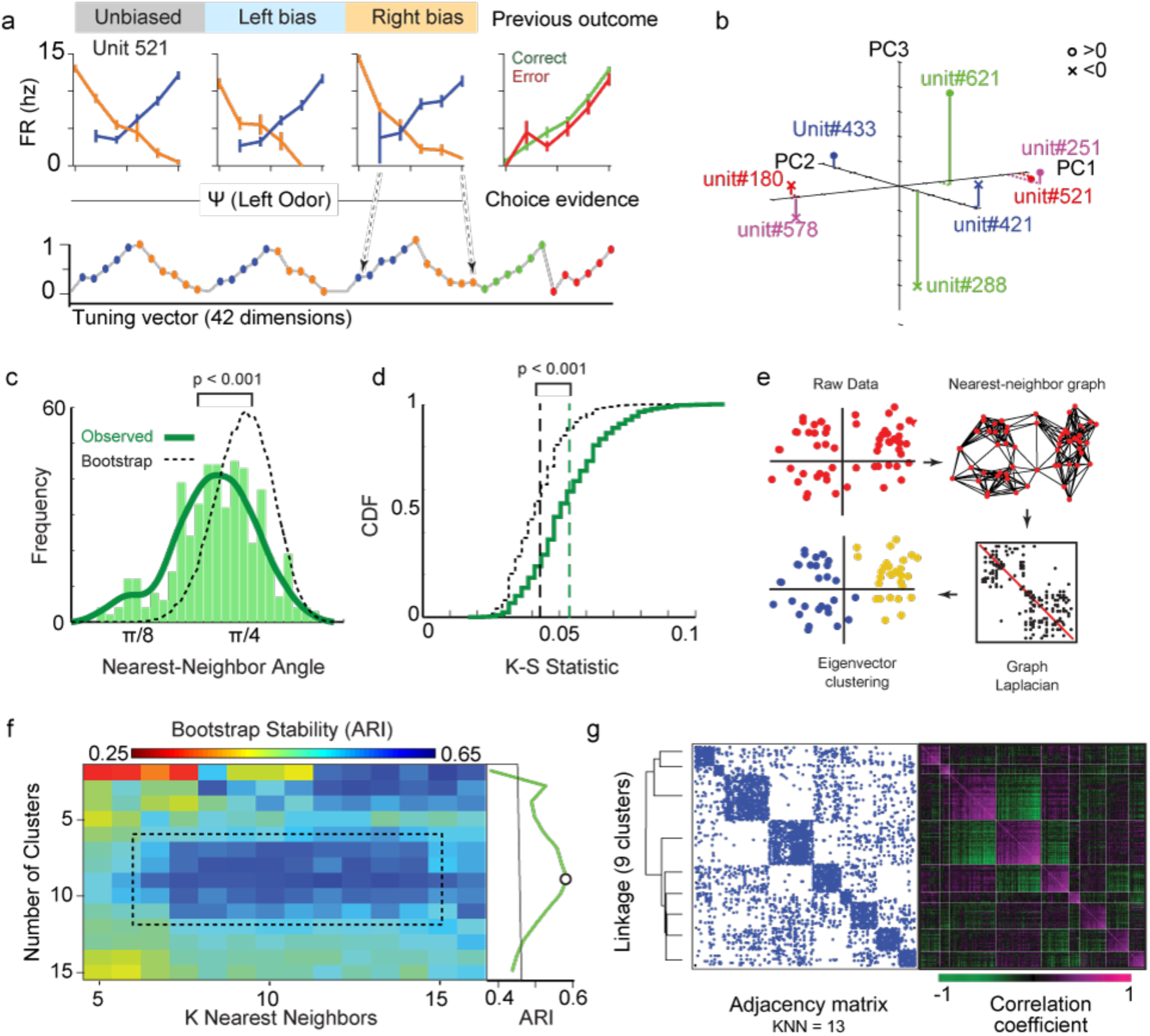
OFC neurons can be categorized into 9 discrete clusters. *a*. The response profile of each neuron was generated from 42 behavioral conditions, corresponding to the average firing rate of each neuron during the anticipation window. ***b*** Location of several example cells in the first three dimensions of the resulting eigenspace. ***c-d*** The ePAIRS (c) and eRP (d) tests reveals strong signatures of non-random clustering in the OFC population. For ePAIRS, nearest neighbor angles were smaller than expected, suggestive of clustering (rather than dispersion). ***e*** The analysis procedure for spectral clustering of OFC response profiles. Spectral clustering relies on clustering a connectivity graph built by linking similar neurons, and effectively identifies robust clusters, despite the confounds of an arbitrary topology in high-dimensional space (see Methods). ***f*** Clustering results and hyperparameter selection. A set of 90% bootstrap samples are clustered for each combination of major hyper-parameters (number of nearest neighbors, k; number of clusters, c); each combination of hyperparameters is evaluated for stability using the Adjusted Rand Index (see Methods). For this dataset, clustering results are most stable for 9 clusters, across a wide range of *k* nearest neighbors (maximum at k=13). ***g*** Clustering results. The relationship between the 9 clusters can be examined visually using a linkage map (left), as well as by observation of the structure of the nearest-neighbor graph (center) and the within-cluster and between-cluster correlation coefficient (right).

We next sought to assess whether the OFC population shows *random mixed selectivity,* which is equivalent to saying that the population as a whole is category-free(Raposo et al., 2014). To do so, we developed two novel statistical tests for *random mixed selectivity* that account for the geometry of neural data (ePAIRS and eRP; Raposo et al., 2014). Using these tests, we observed significant deviations from random mixtures – in particular, neurons with similar representations are more similar than expected by chance – suggesting that the population of neuronal representations is likely to show significant clustering (Fig. 3c-d).

Given the non-random structure of the OFC population representation, we next sought to determine whether the tuning profiles of OFC neurons could be clustered into a set “categorical” representations. To do so, we looked for groups of neurons with similar response profiles using spectral clustering, a non-linear clustering technique that exploits nearest-neighbor distances. In this context, spectral clustering relies on building a neighborhood graph in which neurons with very similar responses are connected; clusters are then identified as strongly connected subcomponents of this graph (Fig. 3e; **Methods**). This technique mitigates noise, allows for arbitrary cluster number and shape, and avoids the problems of distance measurement in high dimensions. To maximize the reliability of this result, we used bootstrap cross validation to evaluate the cluster stability across changes in two critical hyperparameters (the number of nearest-neighbors **k**, and the number of clusters **c**) using the Adjusted Rand Index (ARI; see **Methods**).

When applied to the OFC population, this method reveals a reliable segmentation into 9 clusters of neurons, each encoding a distinct and categorical representation (Fig. 3f). Visual examination reveals strong structure in the resulting network graph, as well as in the cluster correlation matrix (Fig. 3g). The number of clusters was set by cross-validation and is insensitive to variation in *k* across the range in which the neighborhood graph is unconnected (*k = 5*) to densely connected (*k* > 15). In addition, the observed peak of ARI was significantly higher than expected of random or trial-shuffled datasets (ARI ∼0.65; *p* < 0.01; *t*-test). These results were not affected by missing-data imputation (Extended Data Fig. 6), and did not arise from a simple segmentation of neurons from different animals (Extended Data Fig. 5c; **Supplemental Table 1**). Overall, neuronal response profiles are strongly correlated within clusters, but uncorrelated or negatively correlated between clusters (Fig. 3g).

We note that there are multiple ways in which we were biased against finding such clusters. First, any spike sorting errors that contaminate single units would undermine cluster divisions. Second, our behavioral task is insufficient to probe every behavioral contingency that could differentiate between otherwise similar neurons. Third, sampling limitations limit both the number of neurons recorded and the accuracy of each response profile, constraining any clustering result. Lastly, we note that the clustering and cross-validation procedure itself is biased against small clusters (**Methods**). For all these reasons we expect our results to provide an underestimate of how structured OFC representations may actually be.

To examine each response type more closely, we first generated average response vectors for each cluster. Surprisingly, although these 9 clusters were identified entirely without model input, the average tuning profile of each cluster resembles a putative decision-variable. These include familiar representations that can be directly compared to the exemplar neurons described above: *reward size* (Cluster H), positive and negative confidence (*confidence*^[-]^, cluster A; *confidence*^[+]^, Cluster B), and negative integrated value (*value*^[-]^, Cluster C). For each of these clusters, the average tuning curves closely match the variables present in the decision-variable model (Fig. 1f). We also observed other important variables, including responses associated with choice (*choice side*^[L]^, Cluster D; *choice side*^[R]^, Cluster E); the average reward size within a given block (*average reward size*, Cluster G), and the outcome of previous trials (*previous outcome,* Cluster F). In addition, we note several subtle deviations from our expectations, including the apparent opponency of many but not all clusters, and the somewhat idiosyncratic representation found in the smallest cluster (Cluster I).

We took two approaches to quantitatively assess how well these 9 categorical OFC responses types correspond to specific decision variables. First, we analyzed the response of neurons in two major clusters, corresponding to *confidence* and *integrated value*, variables for which we can make a number of additional predictions. Second, we asked more generally how well the clusters correspond to a decision-variable model of our task.

We first analyzed the cluster encoding a putative representation of confidence^(+)^ (Cluster B). For both averaged and individual neurons, we observed activity proportional to the odor evidence supporting each choice (Fig. 5a.i-iii, viii). In addition, this cluster conforms to three predictions of the computation of statistical decision confidence(Kepecs et al., 2008). First, activity decreases with stimulus difficulty for correct but increases for error choices. Second, the relationship between choice accuracy and stimulus contrast depends in firing rate, suggesting an encoding of subjective difficulty rather than stimulus type (Fig. 5a.v). Third, firing rates predict choice accuracy regardless of stimulus (Fig. 5a.vi). Similar responses were observed for the cluster encoding negative confidence, with appropriate sign reversals (Cluster A; Extended Data Fig. 7).

We next analyzed Cluster C, which resembles a decision-variable for negative integrated value (*value*^[-]^; Fig. 5b). Since integrated value is defined as the probability of reward for a given choice (decision confidence) multiplied by reward size, we tested for the influence of these representations. The average response profile of Cluster C reflects this integration, revealing signatures of both variables seen in average activity (Fig 5b.i-ii) and strong correlations with model-based estimates of integrated value (Fig. 5b.iii, Fig. 5b.v). Importantly, the firing rate of Cluster B neurons tracks behavioral accuracy below 50% – reflecting outcome probability for decisions made under reward bias, when animals do not always chose the maximum confidence option (Fig. 5b.iv-vi).

These results show that the largest clusters of OFC neurons align closely to two critical components of the decision-variable model of OFC function: confidence and value. Consistent with the idea that individual OFC neurons are highly biased for coding coherent representations of integrated value (Fig. 5c), in spite of the anti-correlation between the two variables (correlation coefficient: -0.26), we did not observe other clusters encoding the incoherent combinations of confidence[+/-] and reward size[-/+] (Fig. 4).

**Figure 4.**
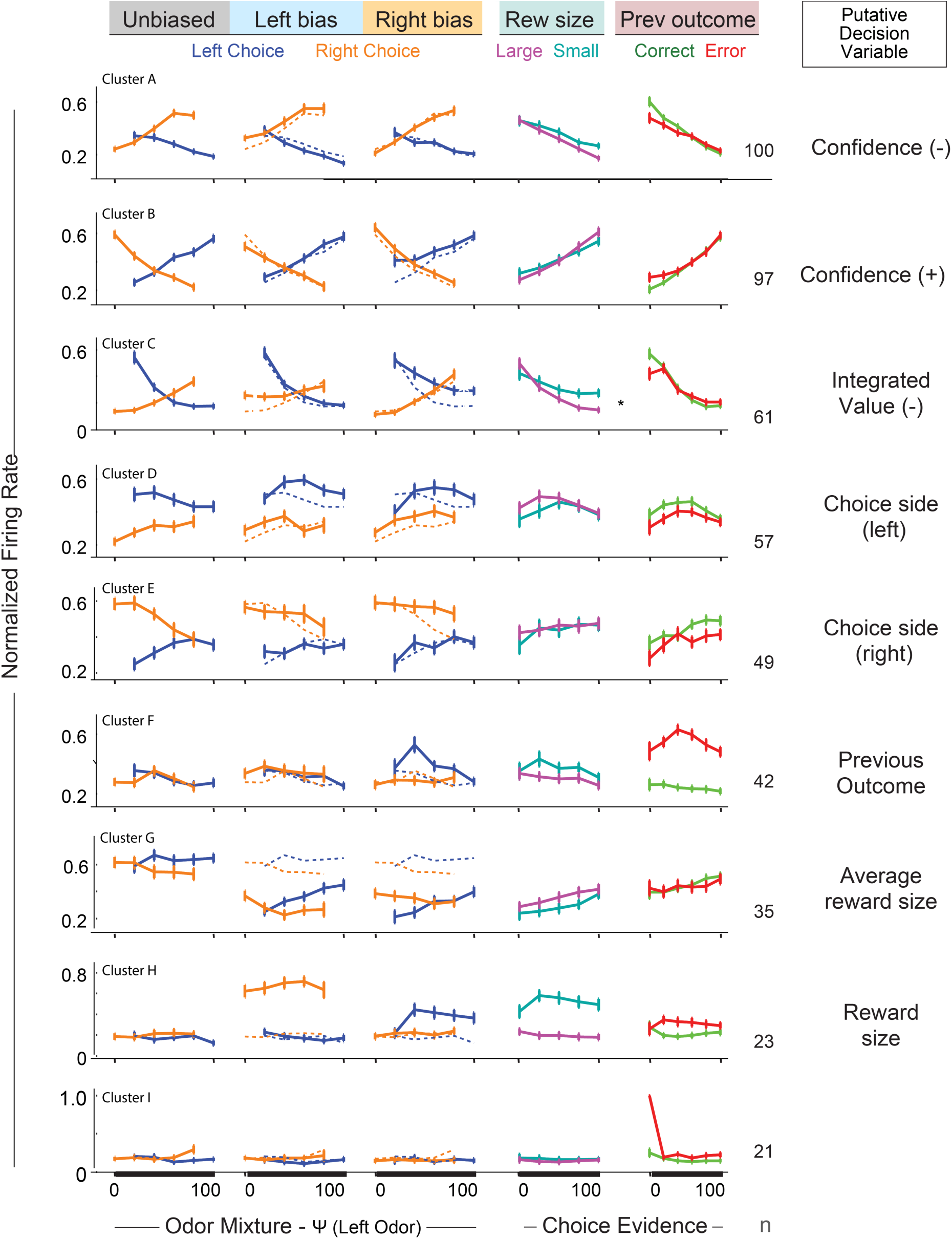
Identified Clusters of OFC Neurons. *a-j*. Average response profiles of each of the 9 identified response clusters. For each cluster, the normalized firing rate is shown for all 42 behavioral conditions used to generate the clustering results (responses conditioned on stimulus and choice, *Unbiased*, *Left Bias*, and *Right Bias* blocks; conditioned on outcome of the previous choice and the evidence supporting the current choice, *Prev Outcome*). In addition, normalized firing rates are shown conditioned on the size of the the reward associated with the choice port (*Rew Size*). For each cluster, we also note the corresponding putative decision variable.

To generalize this approach to all decision variables we turned to an alternative approach, and asked how well the representations encoded individual OFC neurons could be described using a decision-variable model. To do so, we generated a model carrying "elementary task variables" such as the *odor stimulus*, *choice side*, and *expected reward*; more complex representations of "canonical decision variables" such as *confidence* and *integrated value* arise from the combination of a small number of these variables. Confirming the idea that this model is a reasonable starting point to assess OFC activity, we observed that individual OFC tuning curves were well fit by this model, with significantly lower error than observed for fits of trial-shuffled neurons (*p* < 1E-64, Fig. 5d).

**Figure 5.**
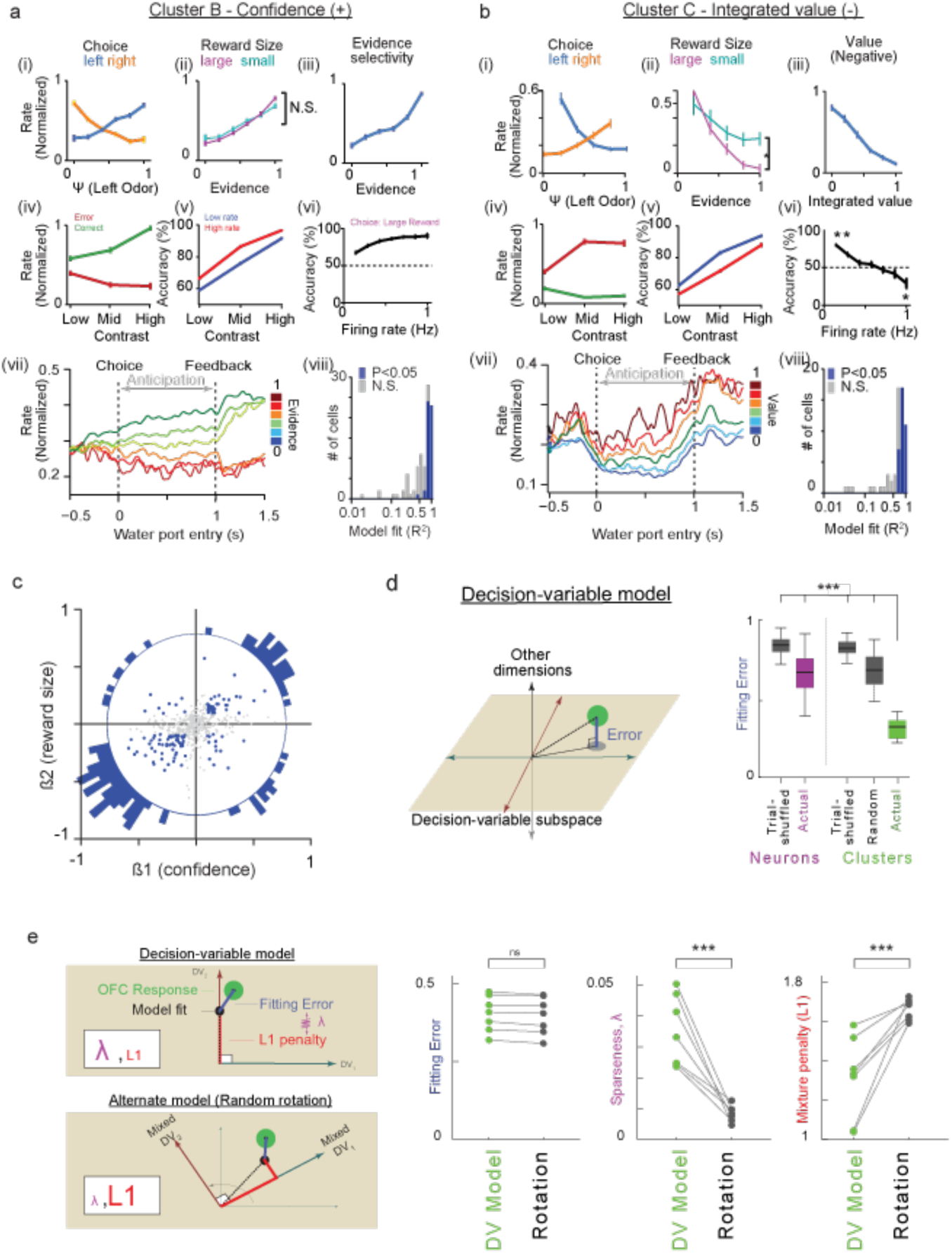
Analysis of OFC Clusters. a. Characteristic response profile of **Cluster B**, corresponding to a decision-variable representation of *Confidence (+)*. Averaged normalized responses for neurons in this cluster are shown for various conditions: ***i*** stimulus and choice, ***ii*** expected reward size, ***iii*** overall choice evidence selectivity, **iv** stimulus contrast, ***v*** choice accuracy as a function of contrast, *and* ***vi*** choice accuracy as a function of firing rate. ***vii*** Peri-stimulus time histogram of normalized FR, conditioned on the evidence supporting choice. ***viii*** R2 of each cells in the cluster based on a trial-by-trial fit to choice evidence. **b** Characteristic response profile of **Cluster C**, corresponding to a decision-variable of Integrated Value (-). Sub-panels *i-viii* match those for **Cluster B** (a)**. c** Single-neurons encoding combinations of confidence and reward size show coherent integration. Each neuron’s response profile was fit to a two-parameter model representing confidence and reward size. For most neurons, ß coefficients for each component share the same sign. Data are shown for all neurons (gray) and neurons that show significance in both beta coefficients (blue; p < 0.01 threshold), as well as a polar histogram (p < 0.01 vs. uniformity). **d** The model-free clustering procedure identifies clusters of neurons that are well-represented by a combination of elementary task variables such as stimulus type and choice. Mean response profiles of single neurons and averaged clusters were fit by this model: fitting error was lower for actual neurons than for trail-shuffled control (p < 0.001), and error was dramatically lower for fits of cluster averages than for single neurons, clusters of trial-shuffled neurons, or random clusters. Error distributions were compared under cross-validation by bootstrap test, *** p < 0.001. ***e*** A LASSO analysis of individual clusters reveals that most clusters can be understood as ′canonical′ decision-variables that arise as a sparse mixture of elementary task variables. We used two sets of decision-variable models to fit the average cluster response profile: the canonical model using elementary task variable such as stimulus and choice side, and a suite of alternate models in which these elementary task variables are randomly mixed (eg. choice-side + reward block type – stimulus evidence). Importantly, both models outline the same decision-variable subspace, but represent canonical decision-variables (such as choice value) with very different sparsity. The true DV model generated fits with higher sparseness (λ) and a smaller mixture penalty (L1), suggesting that individual clusters of OFC are well described by the canonical decision-variable model. Sign rank test, *** p < 0.001.

We next considered the variables encoded in each representational cluster. We hypothesized that each cluster corresponds to an unknown decision-variable, with each neuron serving as a noisy exemplar. Under this assumption, we expect that the average response for each cluster will show lower fitting error to the decision-variable model than observed for single neurons. Indeed, we observed that the fitting error of averaged clusters was dramatically reduced compared to single neurons, clusters of trial-shuffled neurons, or clusters of randomly chosen neurons (*p* < 1E-10 for all tests; *t*-test; Fig. 5d). This result suggests that the clustering procedure – which we emphasize is entirely free of model-based assumptions – appears to isolate representations that align closely with the decision-variable model.

However, this approach leaves open the possibility the representation carried in each neuronal cluster corresponds to complicated mixtures of many elementary task variables, rather than the simple mixtures that reflect canonical model-based decision-variables such as confidence and value. To disambiguate these possibilities in the most general way possible, we examined whether the neural clusters were well-represented using a sparse regression that applies a cost for such mixed representations.

Using cross-validated LASSO regression, we found that the OFC cluster representations were well fit by a small number of canonical decision-variables (Fig. 5e). As a control, we also built a library of random models in which elementary task variables are randomly mixed (i.e., the basis set of elementary task variables is randomly rotated). Critically, these non-canonical models allow for representations of canonical decision variables, but these representations are no longer sparse. As expected, the neuronal clusters were well fit to the canonical model, with higher sparsity (*p* < 0.001) and a small mixture penalty (L1 penalty, *p* < 0.00042) than observed for non-canonical models, without an overall increase in error (*p* > 0.05). This analysis reveals that each cluster is best understood as a mixture of elementary task variables, and that these mixtures directly encode the canonical decision variables that populate a normative choice model.

## DISCUSSION

We sought here to characterize the diversity of neuronal representations in orbitofrontal cortex. We used a behavioral task that combines perceptual and value-guided decision making so that we could interrogate OFC representations across numerous dimensions. Our main result is that OFC representations are heterogeneous but highly structured – encoding a small set of categorical representations that correspond to the appropriate decision-variables.

Previous approaches to studying decision-variables in cortical populations have adopted a model-based framework, in which a hypothesized cognitive variable is tested by “decoding” from population-wide activity. Using a single-cell or population decoder approach, such analyses have identified the OFC as a major hub for important internal variables that may support decision-making, including outcome expectancy (Feierstein et al., 2006; Kennerley et al., 2009; McGinty et al., 2016; Morrison and Salzman, 2009; O’Neill and Schultz, 2010; Ogawa et al., 2013; Roesch et al., 2006; Schoenbaum et al., 2003, 1998; Thorpe et al., 1983; Tremblay and Schultz, 1999; Wallis and Miller, 2003), decision confidence (Kepecs et al., 2008), and the value of chosen rewards (Abe and Lee, 2011; Padoa-Schioppa and Assad, 2006; Roitman and Roitman, 2010; Steiner and Redish, 2014; Sul et al., 2010). A major challenge to this approach is the threat of confirmation bias, as it is difficult to compare or reject alternate representations and models.

Here we invert this approach, by using model-free methods to categorize population representations in an unbiased way. We then map these onto model-based decision-variables, and suggest that many hypothesized variables, including choice side, decision confidence, expected reward size, integrated value, and previous outcome, have a privileged representation within OFC. These results thus constrain many models of OFC function and clarify the representational logic of OFC.

Our findings also address a major architectural question: how are representations of important variables shared across neurons? Neurons that mix different variables, such as stimulus type or choice are said to show mixed selectivity. When variables are mixed randomly across neurons, the population is said to be “category free” and show *random mixed selectivity*. Category-free representations may be important for associative learning (Rigotti et al., 2013), and have been reported in both parietal and frontal areas (Machens et al., 2010; Mante et al., 2013; Raposo et al., 2014). On the other hand, internal variables such as decision confidence or value can appear mixed when examined as a function of external variables. This is precisely what we observed in OFC clusters, which can be understood as categorical representations of single canonical decision variables, despite appearing mixed from the perspective of the elementary task variables that are under direct experimental control (Fig. 5). This issue is also closely related to the question of whether important variables are integrated coherently, as has been observed primates (Kennerley et al., 2011, 2009; Padoa-Schioppa and Assad, 2006; Raghuraman and Padoa-Schioppa, 2014) or incoherently, as has been observed previously in rodents (Roesch et al., 2006). Our data provide a resolution in that we observe that rats coherently integrate reward size and decision confidence into a representation of value, but also maintain separable populations representing of all three variables (Fig. 4, Fig. 5c).

We suggest multiple reasons why our results diverge from some previous reports about mixed cortical representations. First, the complex behavioral task elicits robust OFC responses that may assist in identifying categorical representations, and in separating reward and sensory uncertainty in particular. Second, the significant complexity of the task used here also provides a technical advantage, in that deviations from random mixed selectivity are more obvious in high dimensions. Indeed, understanding frontal cortex may require such diverse and naturalistic task structures. Finally, the structure of neuronal representations may also fundamentally vary between areas.

Category-free representations may be well-suited to enable fast learning of complex relationships (Mante et al., 2013; Rigotti et al., 2013). In contrast, categorical representations may be better-suited to long-term task learning, including cognitive maps of task space (Wilson et al., 2014), and may support the distribution of distinct decision-variables to different downstream targets.

Many sensory cortical areas appear to exploit a similar sparse and over-complete dictionary, used to efficiently re-encode the sensory world (Lewicki and Sejnowski, 2000; Olshausen and Field, 1996)OFC neuronal responses, similar to other cortical regions (Hromádka et al., 2013), are sparse in that most neurons are inactive in most conditions (Extended Data Fig. 8). OFC representations are also sparse in that they only encode a subset of all possible representations – those in the decision-variable subspace (Fig. 5d) and more so in that they largely encode single decision-variables (Fig. 5e). This representation is also overcomplete (they are linearly dependent, in that more categorical representations than required to span the decision-variable subspace), and redundant (many neurons encode each discrete decision-variable). Thus our results demonstrate that OFC representations provide a sparse and over-complete basis of important decision-variables (Olshausen and Field, 1996; Vinje and Gallant, 2000). In summary, we showed that OFC may use a small dictionary of decision-variable representations that is appropriate for sensory and value-based decision-making. We propose that this representational architecture may be a key feature of cortical populations, fundamental to their computational role – in the case of OFC, decomposing both sensation and action into a suite of distinct decision-variables that support optimal behavior.

## Acknowledgements

We would like to thank past and present members of the Kepecs Laboratory for many helpful discussions. J. Sanders and B. Hangya for help with experimental setup; B. Burbach for technical assistance; A. Lak and T. Gouvea for their comments on an earlier version of this paper. This study was funded by the grants from the Klingenstein, Alfred P. Sloan, and Swartz Foundations.

## Author Contributions

J.H. designed and performed the experiments. J.H. and A.V. analyzed data. A.K. designed the experiments and supervised the project. All authors contributed to writing the manuscript.

## Author Information

The authors declare no competing financial interests. Readers are welcome to comment on the online version of the paper. Correspondence and requests for materials should be addressed to A.K.

## METHODS

### Animal subjects

Three male Long Evans rats (250 g at the start of training) were used for the experiment. The animals were pair-housed and maintained on a reverse 12 hr light/dark cycle and tested during their dark period. Food was available ad libitum, and the rats were placed on a liquid restriction schedule with daily body weight monitoring to ensure that body mass remained within 85% of mass before restriction. Rats received water during each behavioral session and ad libitum in the following 30 min in their home cage. All procedures involving animals were carried out in accordance with National Institutes of Health standards and were approved by the Cold Spring Harbor Laboratory Institutional Animal Care and Use Committee.

### Behavior

#### Behavioral apparatus

All behavioral procedures were conducted as previously described(Uchida and Mainen, 2003). Briefly, behavioral setup consisted of a box with a panel containing the three ports equipped with infrared photodiode and phototransistor. Interruption of the infrared photo beam signaled time of entry of rat into the port. Odors were mixed with pure air to produce a 1:10 dilution at a flow rate of 1 l/min using a custom-built olfactometer (Island Motion, Tappan, NY). Delivery of odors and water reinforcement were controlled using computer data acquisition hardware (National Instruments, Austin, TX) and custom software (BControl, C. Brody) written in MATLAB.

#### Reward biased odor discrimination task

Three naïve rats were trained and tested on a reward-biased two-alternative forced choice (2AFC) odor discrimination task as follows. Rats self-initiated each experimental trial by introducing their snout into a central port where odor was delivered. After a variable delay, drawn from a uniform random distribution of 0.2–0.6 s, a binary mixture of two pure odorants, S(+)-2-octanol and R(-)-2-octanol, was delivered at one of 6 concentration ratios (95/5, 70/30, 55/45, 45/55, 30/70, 5/95) in pseudorandom order within a session. Odor pairs were chosen so that they are equally spaced on a psychometric response scale for ensuring that pairs of odors were equated for difficulty. After a variable odor sampling time, rats responded by withdrawing from the central port, which terminated the delivery of odor, and moved to the left or right choice port. Choices were rewarded according to the dominant component of the mixture, that is, at the left port for mixtures A/B > 50/50 and at the right port for A/B < 50/50. For correct trials, water was delivered from gravity-fed reservoirs regulated by solenoid valves 1 s after the subject entered the choice port. Reward amount, determined by valve opening duration, was set to 0.025 ml in control block and calibrated regularly. The reward amount was reduced to 0.3-0.4x of the original amount (0.3 in two rats, and 0.4 in one rat) either right or left in left-biased reward block or right-biased reward block, respectively. Error choices resulted in water omission and were signaled by white noise (Fig. 1).

Several obvious strategies are sub-optimal in this task. The strategy that relies only on reward probability (ignoring reward size block) loses the opportunities to get bigger reward in bias block as the choice gets difficult. On the other hand, the strategy in which an animal biases its choice on the bigger reward size loses the opportunities to get the sure reward in easier trials. Randomly mixing these two strategies does not increase performance. The optimal strategy in this task is to integrate decision confidence and reward expectation, with maximum bias towards large rewards when decision confidence is low.

For each session, expected reward amount was calculated as

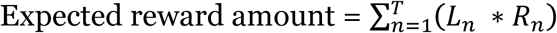

where *L*_*n*_ is reward likelihood in *n*th trial and *R*_*n*_ is reward size in *n*th trial. Reward likelihood was approximated based on the psychometric function across sessions. *T* is actual trial number animals conducted in the particular session. For the strategy without caring reward size (model1) was calculated by setting *R*_*n*_ = 0.025ml in control block and *R*_*n*_= mean reward size for biased block. For the strategy in which animals ignore stimulus evidence (model2), expected reward amount was calculated by setting L_*n*_=0.5 and *R*_*n*_=0.025 ml.

#### Training

In order to be able to perform the task described above, rats went through gradual step by step training procedure, which last typically 4-6 weeks. Initial steps of training consist of imperative trials in which animal is supposed to poke into the central port and collect the water reward subsequently from either of the two reward ports. Choice trials were gradually introduced as the training proceeds; starting with easy choices between pure odor stimuli and advancing gradually with more difficult trials in which odor mixture close to decision boundary is introduced to the animal. During initial training sessions, reward immediately followed the correct choice; later sessions incorporated a delay up to 1sec. Reward bias was introduced gradually, with the smaller reward limited to 30-40% of normal reward amount.

### Surgery

Anesthesia was induced with inhalation of ∼%2.5 isoflurane and retained with intraperitoneal injections of ketamine (50 mg/kg) and Medetomidine (0.4 mg/kg) at the onset of the surgery and supplemented if deemed necessary based on hind leg reflex. Body temperature was maintained using a heating pad (HoMedics, Commerce Township, MI). Rats were stereotactically implanted with custom-made microdrives in the left orbitofrontal cortex (targeted 1.5 mm above OFC (AP+3.7, ML+/-3.2, DV+3.0). Extracellular recordings were obtained with eight independently movable tetrodes using the Cheetah system (Neuralynx) and single units were isolated by manually clustering spike features with MClust (A. D. Redish). Following the surgery, rats were administered ketoprofen (Fort Dodge Animal Health, IA) as an analgesic (5 mg/kg). After 7-10 days of recovery after surgery, action potentials were recorded extracellularly (sampled at 32 kHz) with a Cheetah32 system (Neuralynx, Inc.). During that period, tetrodes were gradually lowered to reach OFC (electrode placements were estimated by depth and later confirmed with histology).

## Electrophysiological recordings

Extracellular recordings were obtained using eight tetrodes. Individual tetrodes consisted of four twisted polyimide-coated nichrome wires (H.P. Reid, Inc., Palm Coast, FL; single wire diameter 12 μm, gold plated to 0.25–0.5 MΩ impedance). Single unit activity was first amplified using a unity-gain op-amp preamplifier fitted to the microdrive array and connected to a bank of eight channel programmable amplifiers (Cheetah acquisition system, Neuralynx, Tucson AZ) via flexible fine wire cables. Signals were band-pass filtered from 0.6-6 kHz and sampled at 31.25 kHz. Individual threshold values were set for each channel such that anytime a recorded voltage crossed the threshold of at least one channel of a tetrode, 32 points were recorded for each of the four tetrode channels. The Cheetah system was also conFigured to record behavioral events generated by the behavioral control computer, as well as the photo beam signals from the odor and goal ports. The tetrode was moved daily (approximately 90 μm) using a microdrive after recording sessions so as to sample an independent population of cells across sessions and all the recorded neurons were analyzed.

### Histology

Once experiments were complete, rats were anesthetized (sodium pentobarbital; overdose) and then transcardially perfused with saline and 4% paraformaldehyde. The brains were removed and 100 μm serial coronal sections were prepared with a vibratome. Recording sites were marked first by coating electrodes with fluorescent dye (Vybrant® DiI, Invitrogen) or electrolytic lesions were made for each electrode.

### Data analysis

#### Behavior

All the analysis was performed in Matlab®. The daily session for each animal consisted of 821 ± 11 (mean ± SEM) trials. Behavioral accuracy was defined as the percent correct choices over the total number of correct and incorrect choices.

Odor sampling duration (OSD) was calculated as the difference between odor valve actuation until odor port exit, with 100 ms subtracted to account for the delay from valve opening to odor reaching the nose(Feierstein et al., 2006) (Fig. 1d). We excluded from calculation of performance accuracy and OSD trials in which odor port withdrawal occurred less than 100 ms after odor onset (0.47 % of all trials) and trials in which no choice was made or choice port entry occurred after the response deadline (0.82% of trials). The psychometric function in Fig. 1C was fitted with a Weibull function using a maximum likelihood method using a linear regression. Curves in Fig. 1e were fit with a following the 4 parameter nonlinear regression model.

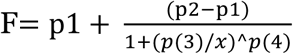

Where x is trial number from block transition and p1 to p4 are fit values using least squares estimation. Error bars are mean ± s.e.m. (n across rats) or mean ± s.d. (n across sessions). The effect of odor contrast on accuracy or OSD was tested using one-way ANOVA with pairwise comparisons between different mixture contrast ratios (*MULTCOMPARE* function in Matlab®) at a significance level of *P* < 0.0125 (i.e. adjusted for multiple comparisons).

#### Integration model

We aimed to predict choice behavior under reward biased block by the model. We follow four assumptions. (1) Animals expect a constant reward size for each choice option in each reward size block. We ignore the update of reward size expectation which occurs in block transitions for simplicity. (2) Animals have estimation of likelihood of outcome (correct or error) for each choice option (left or right) once they receive odor mixture in each trial based on signal to noise ratio of odor stimulus mixture(Kepecs et al., 2008). (3) Those expected values are integrated in multiplicative way for each option. (4) Animals choose the option with larger value. This leads to the choice probability of one of options is proportional to the relative value of the choice option.

Expected relative reward size (R_A_) for each option in a particular block is as follows.

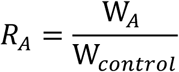

Where W_A_ is water reward amount for choice option A (i.e., left or right) in reward biased blocks and W_A_ is water reward amount in control block (W_control_). Likelihood (L_X_) of outcome for choice X for a given stimulus is directly estimated by the psychometric response function when reward size is equal (control block). For example, if animals chose 98% of left side in response to odor pair [95/5%, A/B odor pair], animals’ estimated likelihood of chosen outcome for the odor pair would be 98%. Likewise, if probability of left choice to the odor pair [5/95%, A/B] is 3%, the estimated likelihood of outcome for the odor pair is 3%. However, because we chose odor pairs so that they maximally utilize the psychometric response space in the first place (linear line of psychometric response to 6 odor pairs, Fig. 1b), we simply approximated the outcome likelihoods for left choice as [0, 0.2, 0.4, 0.6, 0.8, 1] for 6 sets of left odors as the odor ratio in the mixture that supports left choice increases. The outcome likelihoods for right choice equal to the inverse of outcome likelihoods for the left choice (i.e., [1, 0.8, 0.6, 0.4, 0.2, 0]).

The choice value for choice option X (left or right) was estimated by the following function.

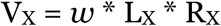

Where *w* is the weighting coefficient determined by fitting the choice behavior as described below (*w* = 1 for unmanipulated side).

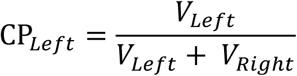

Where CP_Left_ is a choice probability for left choice option. The choice probability function was fitted to the actual behavioral choice probability function (Fig. 1g). Optimal parameter (*w*) was found using a downhill simplex method, *FMINSEARCH* function in Matlab® and coefficient of determination R^^2^^ was calculated by squaring correlation coefficient between model with the determined parameter and the actual choice data. The chosen value to evaluate neuronal representations was estimated by a following function.

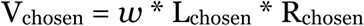

Where *w* from the behavioral fitting above was used and L_chosen_ was called “choice evidence”

We extended this model to explain choice behavior during control blocks, under the assumption that that animals have the trial-by-trial expected reward value based on the outcome of a previous trial integrating with the likelihood for odor categorization (Extended Data Fig. 2).

#### Electrophysiology

##### Spike detection and sorting

Spikes were manually sorted into clusters (presumptive neurons) off-line based on peak amplitude and waveform energy using MClust software (A.D. Redish). Clusters were considered as single units only when the following criteria were met: (1) refractory period violations were less than 1% of all spikes and (2) the isolation distance, which was estimated as the distance from the center of identified cluster to the nearest cluster on the basis of the Mahalanobis distance, was more than 20.

##### Spike train analysis

We analyzed single unit data from 68 sessions for each animal during the performance of the spatial discrimination task. Unless otherwise stated, spike trains were smoothed by convolution with a Gaussian kernel (σ = 15 ms) to obtain a spike density function (SDF) for the analysis of the temporal profile of neuronal activity. We focused our analysis on the ‘reward anticipation period’ while rats remained at one of the choice ports.

Neuronal response profiles were generated for 485 neurons, including all well-isolated neurons that had a non-zero firing rate during the anticipation window. Response profiles were generated from 48 conditions (Fig. 3) with six conditions dropped due to frequent missing values (Extended Data Fig. 6) to leave 42 conditions.

The matrix of response profiles were subject to denoising and imputation of sparse missing values using probabilistic PCA (pPCA). Eigenvectors of this decomposition are shown in **Extended Data Figure 5**), of which approximately the first 7 (accounting for ∼60% of observed variance) resemble somewhat distorted mixtures of common decision-variables.

The interpretability of such eigenvectors as dominant neuronal tuning curves is limited by the assumption that the data are distributed as a multivariate Gaussian. As a simple example, we note that the number of eigenvectors resembling decision-variables can be smaller than the actual number of unique decision-variables that are represented, because decision-variables are not linearly independent. For example, a 2D space whose basis vectors represent confidence and reward size could contain representations of three distinct decision-variables (confidence, reward size, and choice value). We therefore sought to understand whether neurons in such a space show random mixed selectivity, or instead form categorical representations that align to putative decision-variables.

To retain full population diversity in downstream analyses, coefficients were truncated to retain 90% of between-neuron variance, resulting in a set of coefficients for each neuron in a 21-dimensional response space. We next sought to understand how the population of neurons were distributed in this response space.

Random mixed selectivity in this population was assessed using ePAIRS and eRP. Briefly, these non-parametric tests assess whether the set of neuronal response profile are evenly distributed throughout the representational space, while accounting for differences in dimension size and tolerating changes in the magnitude of each response.

The difficulty of clustering in high dimensions is well known, with many problems arising from the difficulty of estimating large distances or weak correlations reliably. One method commonly used in the machine learning community for examining the structure of a set of points embedded on a high-dimensional manifold is spectral clustering. This technique transforms data from a metric space into a reduced adjacency graph, in which only neuronal response profiles with the strongest correlations are linked, and then attempts to segment or cut this graph into appropriate subcomponents. One intuition is that (normalized cut) spectral clustering identifies a set of cuts in this graph such that a random walk would rarely transition between sub-components(Meila and Shi, 2001). This approach has the advantage of only acting on estimates of small distances (which are reliable even in high dimensions), of being insensitive to whatever manifold that the data may be embedded within, and of being insensitive to the magnitude of each response profile.

Spectral Clustering was performed standard techniques, including Shi and Malik normalization of the Laplacian followed by k-means clustering of the small eigenvectors. For a set of **n p-**dimensional response profiles **X(n,p)**, we first generate an adjacency matrix **A(n,n)**. The **m**th row of A is zero vector supplemented with **k** unit values denoting the **k** rows of **X** that are most similar to row **m** under a cosine (or other) metric. This adjacency matrix is then rendered symmetric as A = A □ A′, and the Laplacian generated as **L = D – A**, where **D** is the degree matrix whose diagonals denote the number of non-zeros in each row of **A**. For the normalized cut operation, this is normalized as **L = (1/D) * L**, where the division **1/D** is taken element-wise (Shi and Malik, 2000). We then find the small eigenvectors of **L** – if these have zero eigenvalues, they represent distinct subcomponents of the adjacency graph. For numerical stability, we treat **c+1** small eigenvectors of **L** as a set of coordinates (c+1 coordinates for each data point), and perform k-means clustering to identify **c** clusters.

The adjacency matrix was generated by identifying the k-nearest neighbors under a cosine or correlation distance metric. Hyperparameters, including number of nearest neighbors (k), the number of clusters (c), were selected by maximizing the adjusted Rand Index (ARI). For ARI analysis, 100 bootstrap samples were generated for each hyper-parameter combination, with a sub-sampling fraction of 0.9. Similar results were observed for both cosine and correlation distance metrics; the results of a cosine metric are shown in Figure 3. We note that k-means is biased against small clusters; although other clustering methods may reveal even greater number of clusters, we chose this approach because it is well validated and in common use.

#### Data availability

The datasets generated during and/or analysed during the current study are available from the corresponding author on reasonable request.

---

**Extended Data Figure 1:**
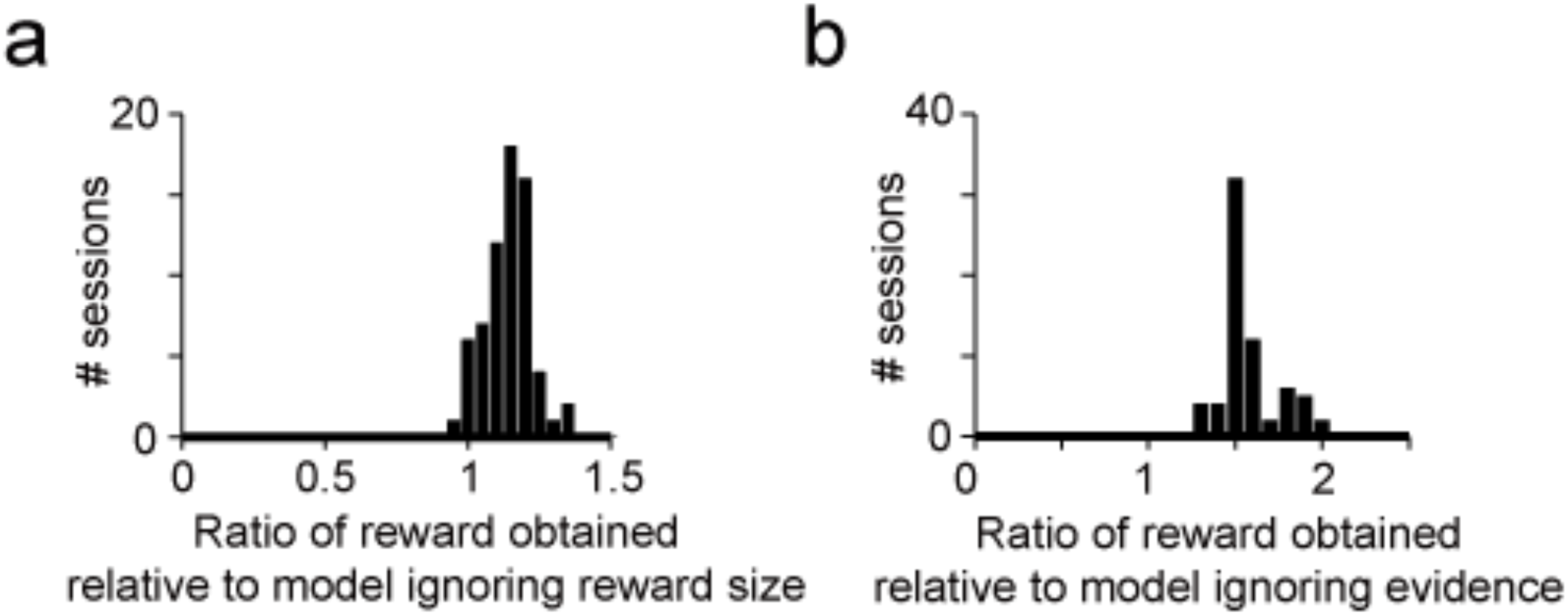
Animals outperform simpler models that ignore either decision confidence or reward size. a. Histogram across sessions, showing the ratio of actual reward to a model relying on odor stimuli but ignoring reward size. **b** Histogram across sessions, showing the ratio of actual reward to a model relying on reward size but ignoring odor stimuli.

**Extended Data Figure 2:**
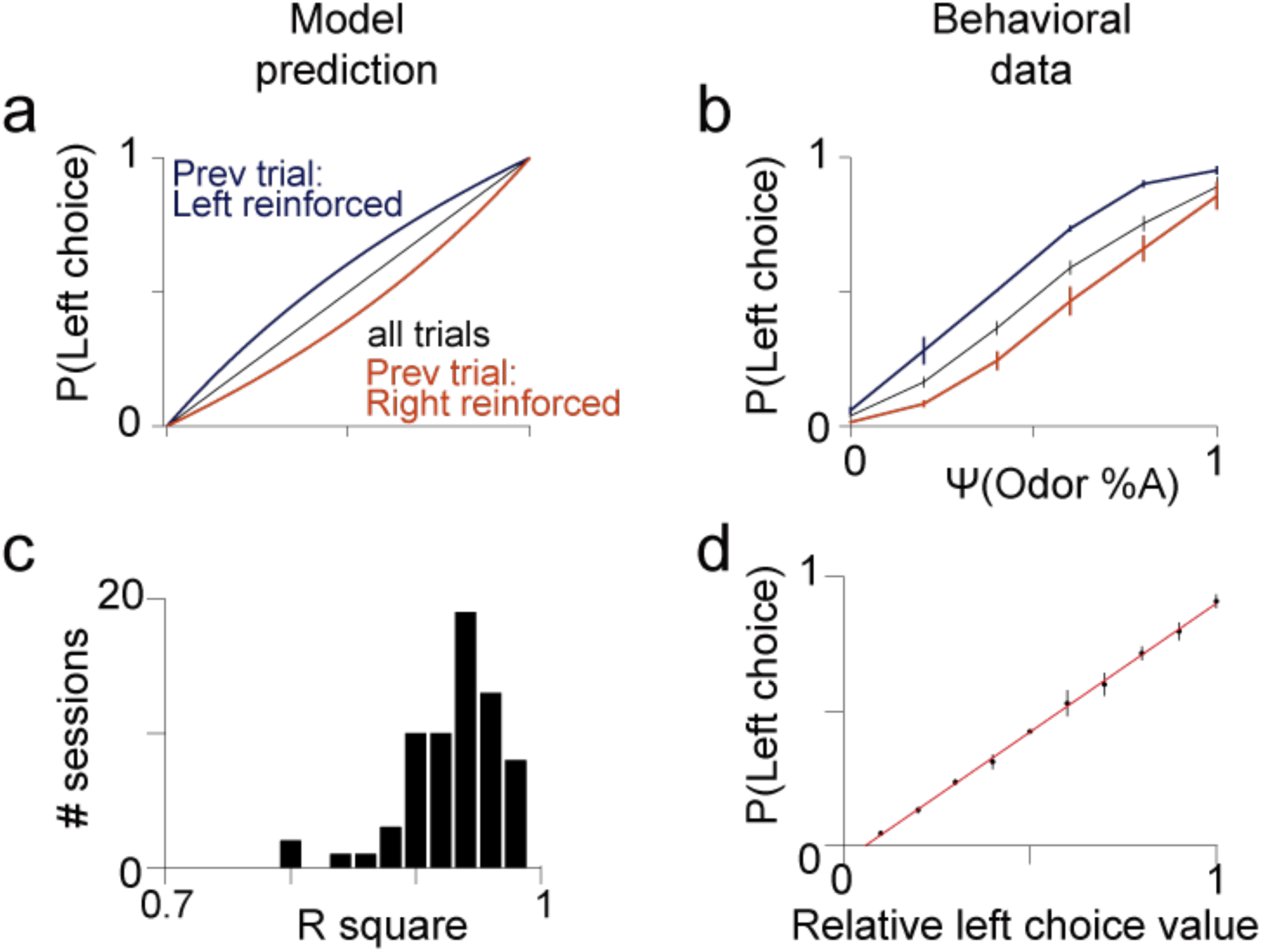
The decision-variable model explains choice bias from previous outcome. a-d. The choice probability for left choice as a function of Ψ(left odor) for all trials or conditioned by left reinforced and right reinforced. Left reinforced indicates that animals are rewarded (correct) in left side in the previous trial or not rewarded (error) in right side regardless of the stimulus conditions used in previous trials. Right reinforced indicates vice versa. The decision-variable model (**a**) accurately predicts changes in choice probability (**b**) arising due to previous outcome. The model provides an accurate fit across sessions (**c**) driven by the relative value of left choice (**d**).

**Extended Data Figure 3:**
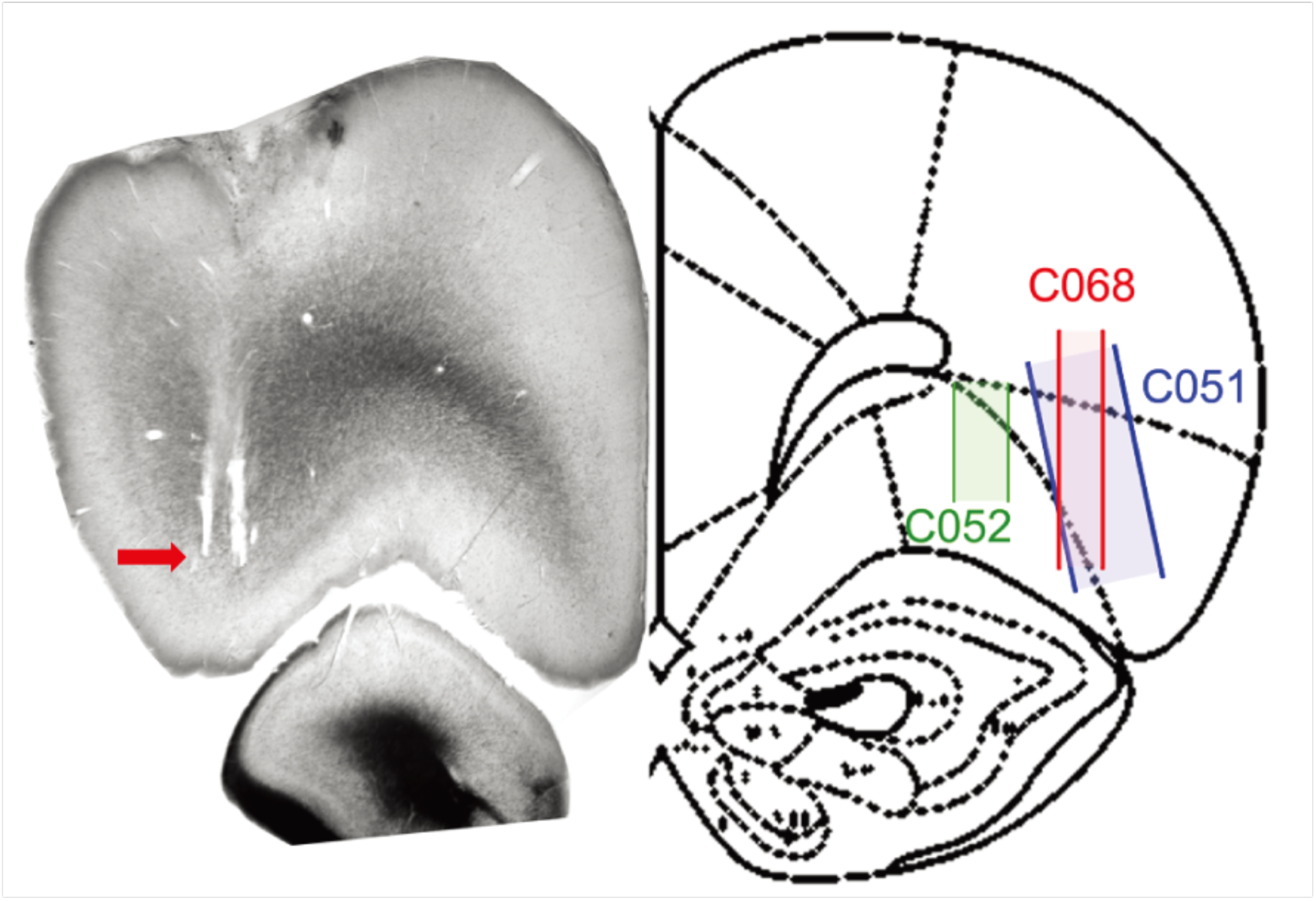
Recording sites in lateral OFC. Recording sites in lateral OFC for all three rats. Histological section shown in left is from C068 as representative, where red arrows indicate the tip of the tetrode bundle. The rectangles in right Figure shows estimated recording ranges where the tetrode bundle was lowered in daily increments.

**Extended Data Figure 4:**
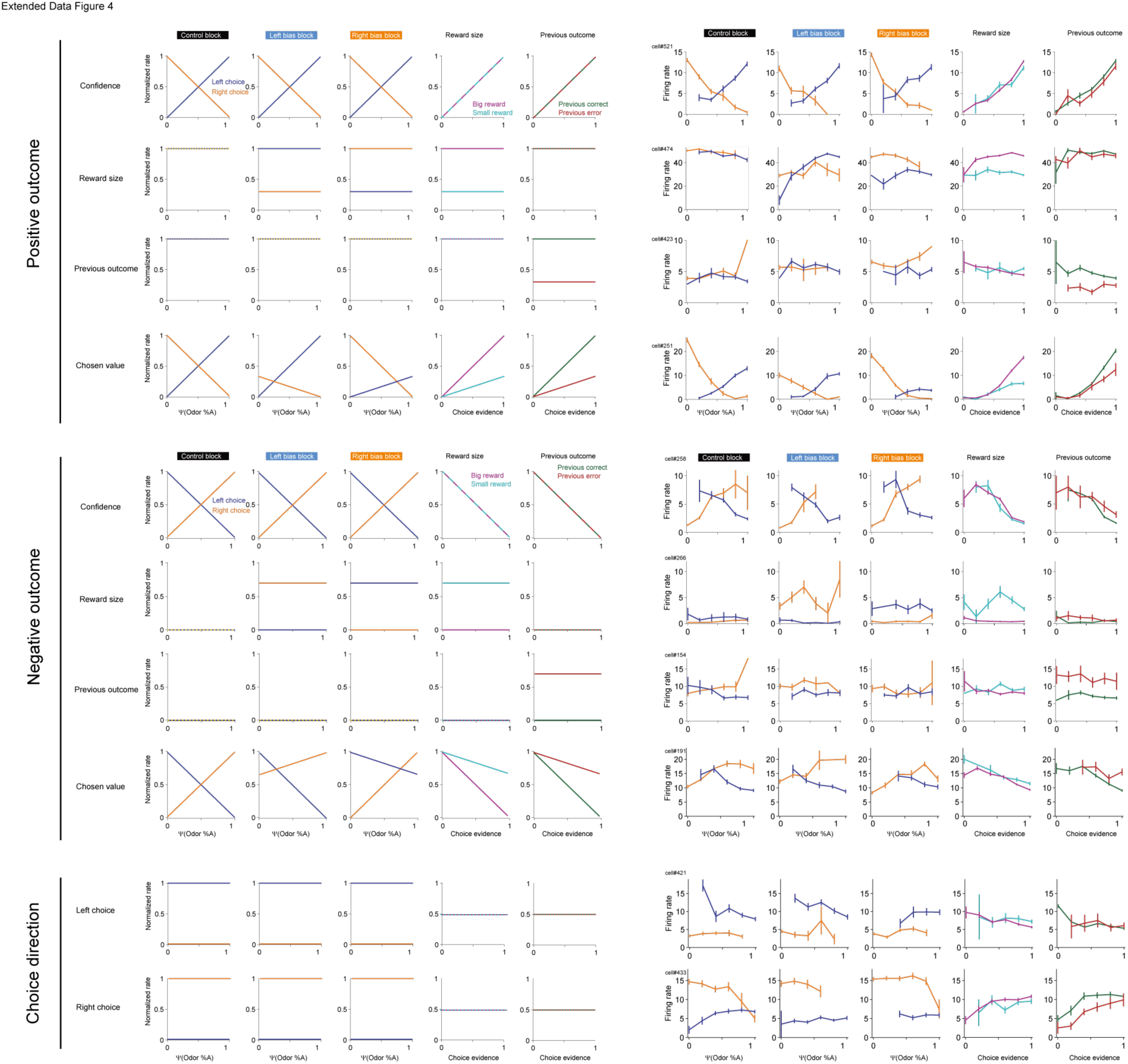
Example response profiles of individual neurons. Example response profiles are shown for several individual neurons. In each case, a schematic tuning curve representing a plausible decision-variable is shown in left panels, while a matching neuronal response profile is shown in right panels.

**Extended Data Figure 5:**
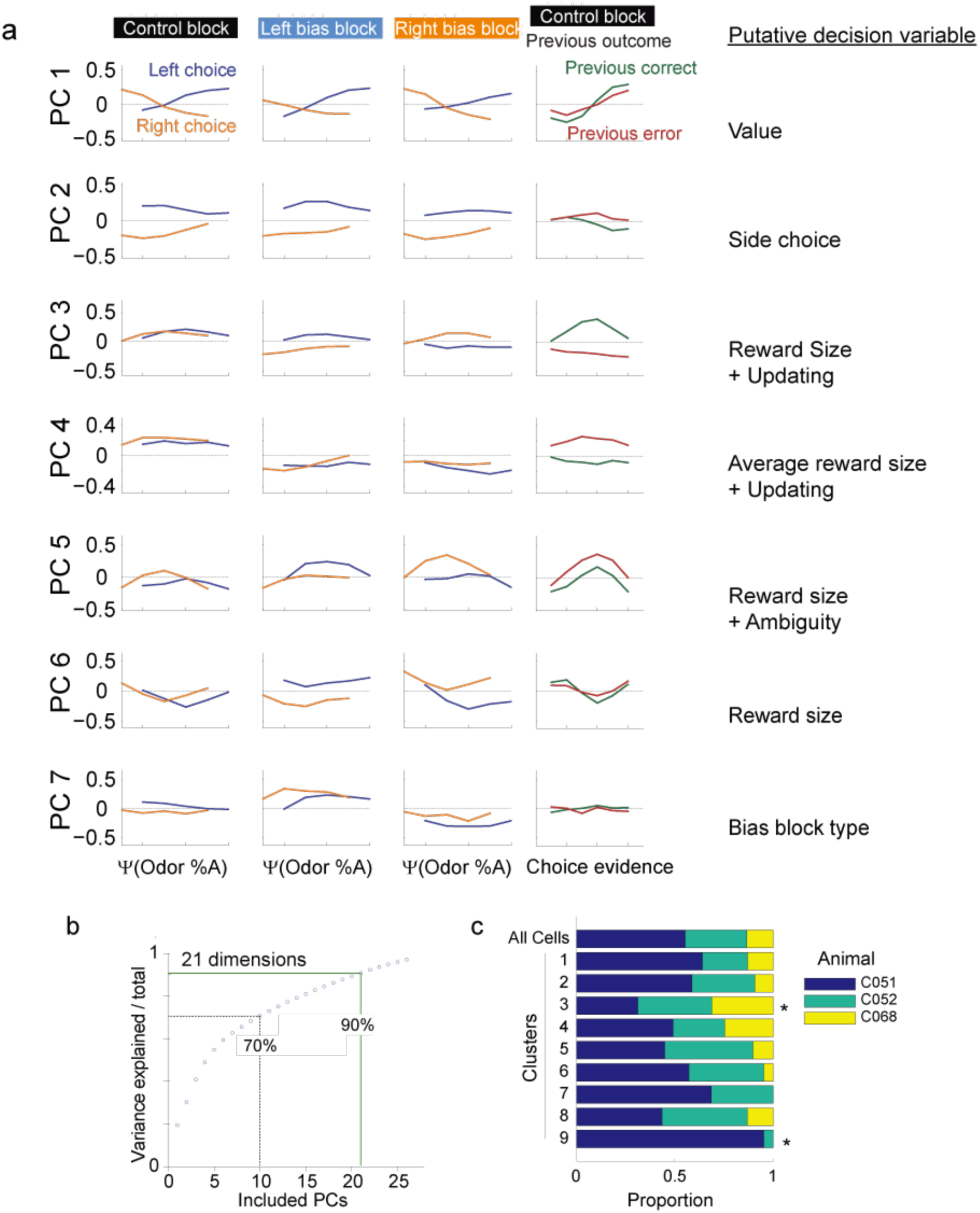
Principal Components Analysis. **a** The 7 dominant principal components arising from a probabilistic PCA decomposition of 485 OFC response profiles, which account for ∼60% of population variance. ***b*** PCA decomposition of the OFC dataset reveals high dimensionality, with 21 principal components required to encode 90% of response profile variability.**c** Clusters are well distributed across animals. Cells in each cluster were generally drawn from all three animals, and rarely showed significant animal-specific bias. *See Supplemental Table 1.*

**Extended Data Figure 6:**
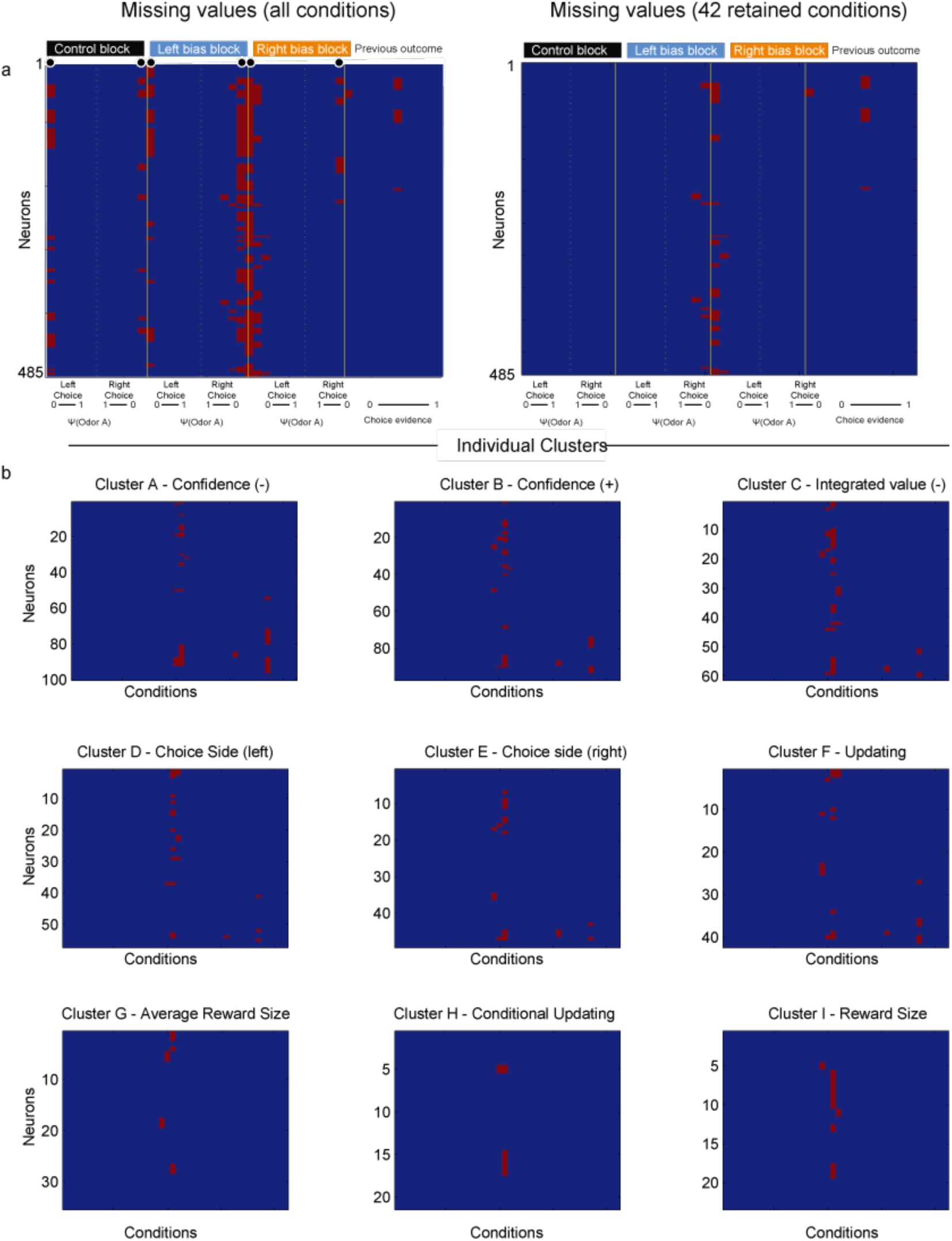
Missing Value Analysis. **a** Missing values are shown for all 485 neuron. *Left* missing values are common for some conditions because animals rarely make errors against high-reliability cues, and are further biased against certain errors during bias blocks. As a result, six possible conditions (black dots) are dropped due to excessive missing data leaving 42 remaining conditions for analysis (right). **b** Missing values are imputed during probabilistic PCA. If these missing values influence clustering, we expect to see a consistent pattern of missing values in certain clusters. Although there is variation in missing values across clusters, there is no obvious pattern of missing data across clusters. **s**

**Extended Data Figure 7:**
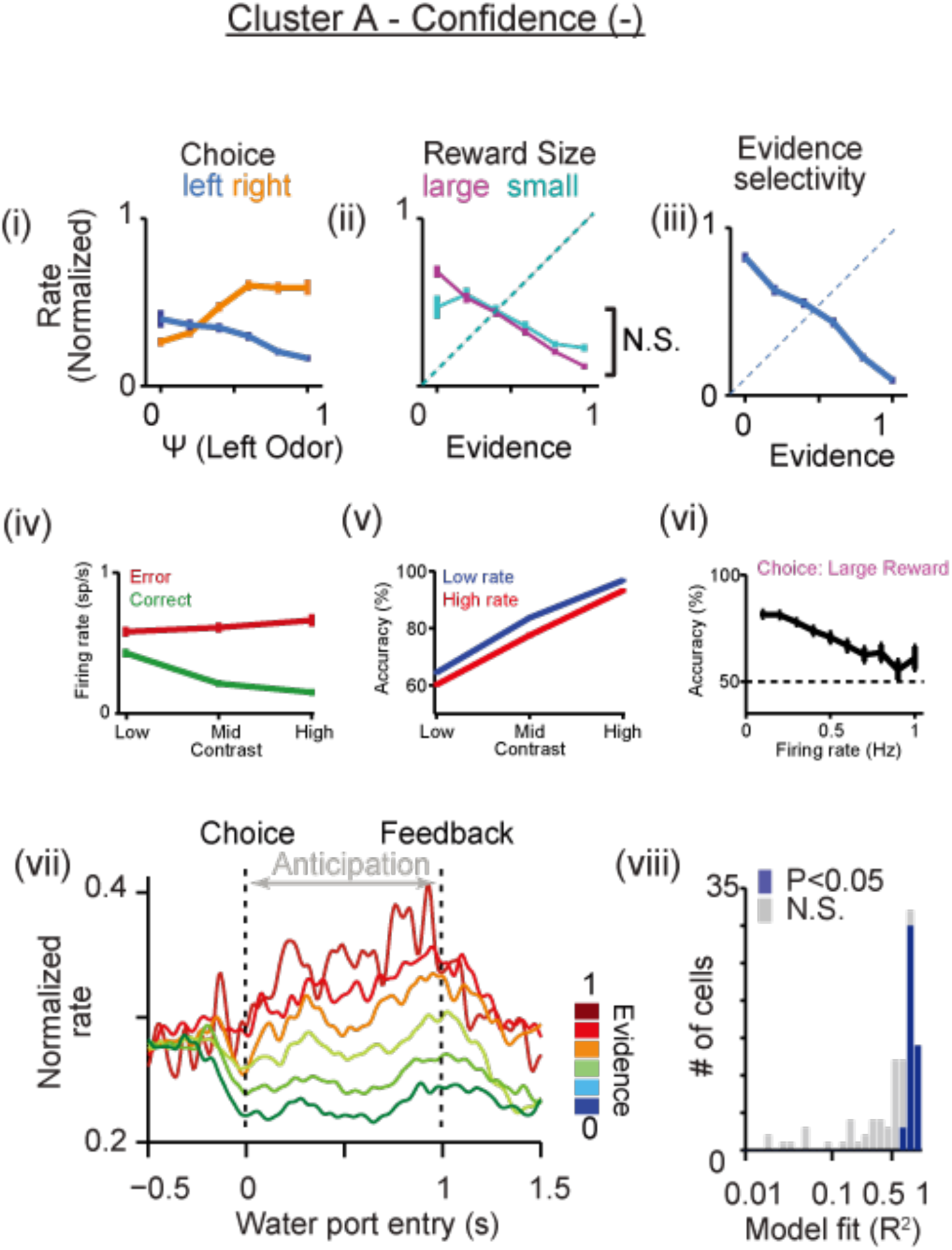
**Characteristic response profile of Cluster A, corresponding to a decision-variable representation of *Confidence (-)*** *Panels match those in* **Figure 5a-b**. Overall, this class of neurons differs from **Cluster B (Figure 5a)** largely in the sign of each relationship.

**Extended Data Figure 8:**
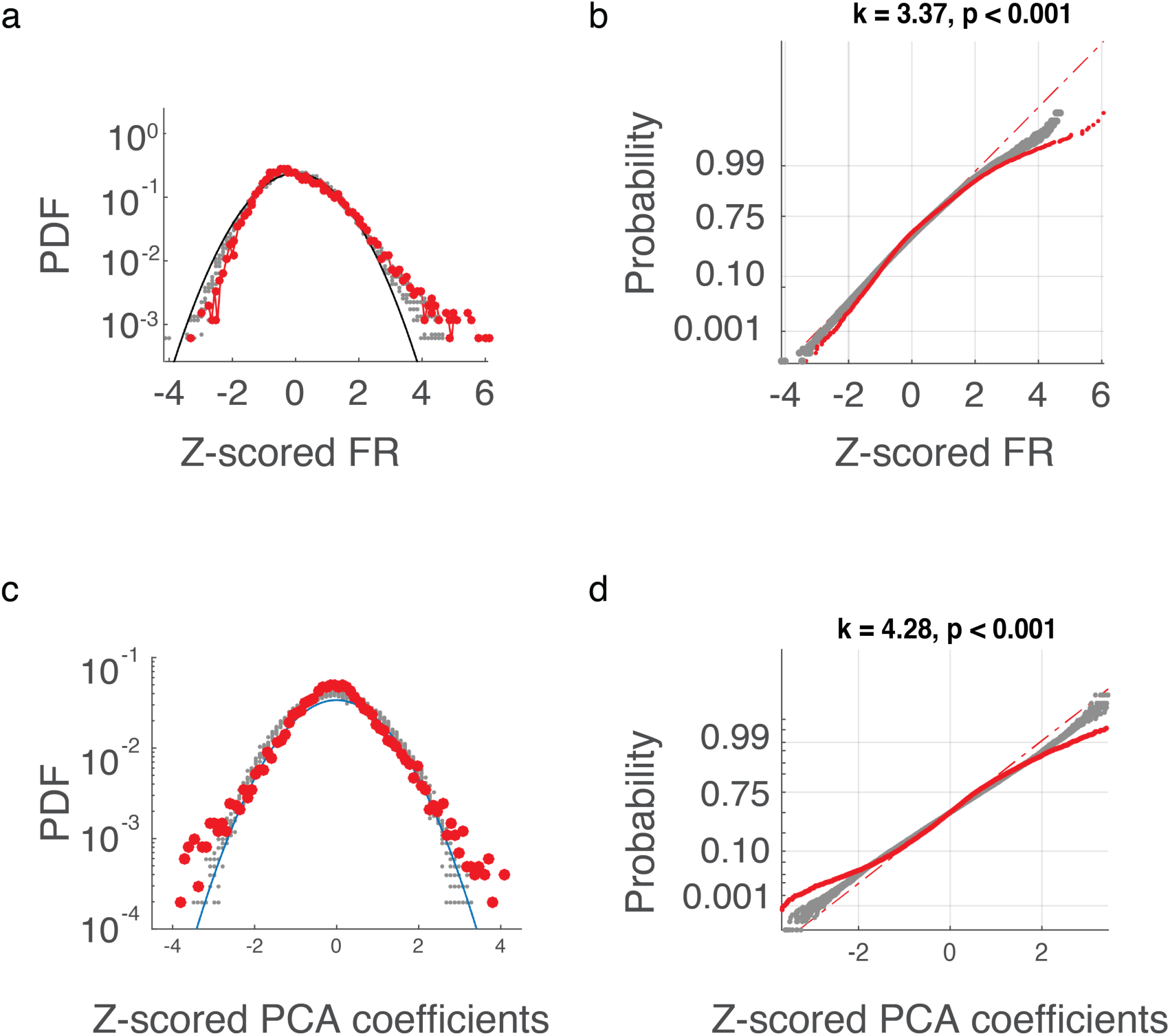
OFC response profiles are sparse by standard measures. **a-b** Firing rates for observed data (red dots) show a right-tailed distribution, with strong activation of most neurons for only a small subset of conditions. This pattern of activation is significantly more sparse than expected from a normal distribution (black line) or trial-shuffled data (gray dots). **c-d** Coefficients arising from PCA analysis show a similar long-tailed distribution compared to a normal distribution (blue line) or trail-shuffled data (gray dots).

## REFERENCES

Abe, H., Lee, D., 2011. Distributed coding of actual and hypothetical outcomes in the orbital and dorsolateral prefrontal cortex. Neuron 70, 731–41. doi:10.1016/j.neuron.2011.03.026

Brody, C.D., 2003. Timing and Neural Encoding of Somatosensory Parametric Working Memory in Macaque Prefrontal Cortex. Cereb. Cortex 13, 1196–1207. doi:10.1093/cercor/bhg100

DiCarlo, J.J., Zoccolan, D., Rust, N.C., 2012. How Does the Brain Solve Visual Object Recognition? Neuron Perspect. 73, 415–434. doi:10.1016/j.neuron.2012.01.010

Farovik, X.A., Place, R.J., Mckenzie, S., Porter, X.B., Munro, C.E., Eichenbaum, H., 2015. Orbitofrontal Cortex Encodes Memories within Value-Based Schemas and Represents Contexts That Guide Memory Retrieval. J Neurosci. 35, 8333–8344. doi:10.1523/JNEUROSCI.0134-15.2015

Feierstein, C.E., Quirk, M.C., Uchida, N., Sosulski, D.L., Mainen, Z.F., 2006. Representation of spatial goals in rat orbitofrontal cortex. Neuron 51, 495–507. doi:10.1016/j.neuron.2006.06.032

Frank, L.M., Brown, E.N., Wilson, M., 2000. Trajectory Encoding in the Hippocampus and Entorhinal Cortex. Neuron 27, 169–178. doi:10.1016/S0896-6273(00)00018-0

Gold, J.I., Shadlen, M.N., 2007. The neural basis of decision making. Annu. Rev. Neurosci. 30, 535–74. doi:10.1146/annurev.neuro.29.051605.113038

Gold, J.I., Shadlen, M.N., 2000. Representation of a perceptual decision in developing oculomotor commands. Nature 404, 390–4. doi:10.1038/35006062

Gutierrez, R., Carmena, J.M., Nicolelis, M. a L., Simon, S. a, 2006. Orbitofrontal ensemble activity monitors licking and distinguishes among natural rewards. J. Neurophysiol. 95, 119–33. doi:10.1152/jn.00467.2005

Hangya, B., Sanders, J.I., Kepecs, A., 2016. A Mathematical Framework for Statistical Decision Confidence. Neural Comput. 28, 1840–58. doi:10.1162/NECO_a_00864

Hromádka, T., Zador, A.M., DeWeese, M.R., 2013. Up states are rare in awake auditory cortex. J. Neurophysiol. 109, 1989–95. doi:10.1152/jn.00600.2012

Kennerley, S.W., Behrens, T.E.J., Wallis, J.D., 2011. Double dissociation of value computations in orbitofrontal and anterior cingulate neurons. Nat. Neurosci. 14, 1581–9. doi:10.1038/nn.2961

Kennerley, S.W., Dahmubed, A.F., Lara, A.H., Wallis, J.D., 2009. Neurons in the frontal lobe encode the value of multiple decision variables. J. Cogn. Neurosci. 21, 1162–78. doi:10.1162/jocn.2009.21100

Kepecs, A., Uchida, N., Zariwala, H.A., Mainen, Z.F., 2008. Neural correlates, computation and behavioural impact of decision confidence. Nature 455, 227–31. doi:10.1038/nature07200

Kiani, R., Cueva, C.J., Reppas, J.B., Newsome, W.T., 2014. Dynamics of neural population responses in prefrontal cortex indicate changes of mind on single trials. Curr. Biol. 24, 1542–1547. doi:10.1016/j.cub.2014.05.049

Lee, D., Seo H.J.M.W., Lee, D., Seo, H., Jung, M.W., 2012. Neural basis of reinforcement learning and decision making. Annu. Rev. Neurosci. 35, 287–308. doi:10.1146/annurev-neuro-062111-150512

Lewicki, M.S., Sejnowski, T.J., 2000. Learning overcomplete representations. Neural Comput. 12, 337–65.

Machens, C.K., Romo, R., Brody, C.D., 2010. Functional, but not anatomical, separation of “what” and “when” in prefrontal cortex. J. Neurosci. 30, 350–60. doi:10.1523/JNEUROSCI.3276-09.2010

Mante, V., Sussillo, D., Shenoy, K. V, Newsome, W.T., 2013. Context-dependent computation by recurrent dynamics in prefrontal cortex. Nature 503, 78–84. doi:10.1038/nature12742

McGinty, V.B., Rangel, A., Newsome, W.T., 2016. Orbitofrontal Cortex Value Signals Depend on Fixation Location during Free Viewing. Neuron 90, 1299–311. doi:10.1016/j.neuron.2016.04.045

Morrison, S.E., Salzman, C.D., 2009. The convergence of information about rewarding and aversive stimuli in single neurons. J. Neurosci. 29, 11471–83. doi:10.1523/JNEUROSCI.1815-09.2009

O'Neill, M., Schultz, W., 2010. Coding of reward risk by orbitofrontal neurons is mostly distinct from coding of reward value. Neuron 68, 789–800. doi:10.1016/j.neuron.2010.09.031

Ogawa, M., van der Meer, M. a a, Esber, G.R., Cerri, D.H., Stalnaker, T. a, Schoenbaum, G., 2013. Risk-responsive orbitofrontal neurons track acquired salience. Neuron 77, 251–8. doi:10.1016/j.neuron.2012.11.006

Olshausen, B.A., Field, D.J., 1996. Emergence of simple-cell receptive field properties by learning a sparse code for natural images. Nature 381, 607–9. doi:10.1038/381607a0

Padoa-Schioppa, C., 2013. Neuronal origins of choice variability in economic decisions. Neuron 80, 1322–36. doi:10.1016/j.neuron.2013.09.013

Padoa-Schioppa, C., Assad, J. a J., 2006. Neurons in the orbitofrontal cortex encode economic value. Nature 441, 223–6. doi:10.1038/nature04676

Pagan, M., Rust, N.C., 2014. Quantifying the signals contained in heterogeneous neural responses and determining their relationships with task performance. J. Neurophysiol. 1584–1598. doi:10.1152/jn.00260.2014

Platt, M.L., Glimcher, P.W., 1999. Neural correlates of decision variables in parietal cortex. Nature 400, 233–8. doi:10.1038/22268

Raghuraman, A.P., Padoa-Schioppa, C., 2014. Integration of multiple determinants in the neuronal computation of economic values. J. Neurosci. 34, 11583–603. doi:10.1523/JNEUROSCI.1235-14.2014

Raposo, D., Kaufman, M.T., Churchland, A.K., 2014. A category-free neural population supports evolving demands during decision-making. Nat. Neurosci. 1–13. doi:10.1038/nn.3865

Rich, E.L., Wallis, J.D., 2016. Decoding subjective decisions from orbitofrontal cortex. Nat Neurosci 19, 973–80. doi:10.1038/nn.4320

Rigotti, M., Barak, O., Warden, M.R., Wang, X.-J., Daw, N.D., Miller, E.K., Fusi, S., 2013. The importance of mixed selectivity in complex cognitive tasks. Nature 1–6. doi:10.1038/nature12160

Rigotti, M., Rubin, D.B.D., Wang, X.-J., Fusi, S., 2010. Internal representation of task rules by recurrent dynamics: the importance of the diversity of neural responses. Front. Comput. Neurosci. 4, 24. doi:10.3389/fncom.2010.00024

Roesch, M.R., Taylor, A.R., Schoenbaum, G., 2006. Encoding of time-discounted rewards in orbitofrontal cortex is independent of value representation. Neuron 51, 509–20. doi:10.1016/j.neuron.2006.06.027

Roitman, J.D., Roitman, M.F., 2010. Risk-preference differentiates orbitofrontal cortex responses to freely chosen reward outcomes. Eur. J. Neurosci. 31, 1492–500. doi:10.1111/j.1460-9568.2010.07169.x

Rushworth, M.F.S., Noonan, M.P., Boorman, E.D., Walton, M.E., Behrens, T.E., 2011. Frontal Cortex and Reward-Guided Learning and Decision-Making. Neuron 70, 1054–1069. doi:10.1016/j.neuron.2011.05.014

Schoenbaum, G., Chiba, a a, Gallagher, M., 1998. Orbitofrontal cortex and basolateral amygdala encode expected outcomes during learning. Nat. Neurosci. 1, 155–9. doi:10.1038/407

Schoenbaum, G., Eichenbaum, H., 1995. Information coding in the rodent prefrontal cortex. II. Ensemble activity in orbitofrontal cortex. J. Neurophysiol. 74, 751–62.

Schoenbaum, G., Setlow, B., Saddoris, M.P., Gallagher, M., 2003. Encoding predicted outcome and acquired value in orbitofrontal cortex during cue sampling depends upon input from basolateral amygdala. Neuron 39, 855–67.

Steiner, A.P., Redish, A.D., 2014. Behavioral and neurophysiological correlates of regret in rat decision-making on a neuroeconomic task. Nat. Neurosci. 17, 995–1002. doi:10.1038/nn.3740

Sul, J.H., Kim, H., Huh, N., Lee, D., Jung, M.W., 2010. Distinct roles of rodent orbitofrontal and medial prefrontal cortex in decision making. Neuron 66, 449–60. doi:10.1016/j.neuron.2010.03.033

Sussillo, D., Abbott, L.F., 2009. Generating Coherent Patterns of Activity from Chaotic Neural Networks. Neuron 63, 544–557. doi:10.1016/j.neuron.2009.07.018

Thorpe, S.J., Rolls, E.T., Maddison, S., 1983. The orbitofrontal cortex: neuronal activity in the behaving monkey. Exp. Brain Res. 49, 93–115.

Tremblay, L., Schultz, W., 1999. Relative reward preference in primate orbitofrontal cortex. Nature 398, 704–8. doi:10.1038/19525

Uchida, N., Mainen, Z.F., 2003. Speed and accuracy of olfactory discrimination in the rat. Nat. Neurosci. 6, 1224–9. doi:10.1038/nn1142

van Duuren, E., Lankelma, J., Pennartz, C.M. a, 2008. Population coding of reward magnitude in the orbitofrontal cortex of the rat. J. Neurosci. 28, 8590–603. doi:10.1523/JNEUROSCI.5549-07.2008

Vinje, W.E., Gallant, J.L., 2000. Sparse Coding and Decorrelation in Primary Visual Cortex During Natural Vision. Science (80-.). 287, 1273–1276. doi:10.1126/science.287.5456.1273

Wallis, J.D., Miller, E.K., 2003. Neuronal activity in primate dorsolateral and orbital prefrontal cortex during performance of a reward preference task. Eur. J. Neurosci. 18, 2069–81. doi:10.1046/j.1460-9568.2003.02922.x

Wilson, R.C.C., Takahashi, Y.K.K., Schoenbaum, G., Niv, Y., 2014. Orbitofrontal Cortex as a Cognitive Map of Task Space. Neuron 81, 267–279. doi:10.1016/j.neuron.2013.11.005

Zariwala, H.A.A., Kepecs, A., Uchida, N., Hirokawa, J., Mainen, Z.F.F., 2013. The limits of deliberation in a perceptual decision task. Neuron 78, 339–51. doi:10.1016/j.neuron.2013.02.010

